# Docking Domain Engineering in a Modular Polyketide Synthase and its Impact on Structure and Function

**DOI:** 10.1101/2023.02.03.526980

**Authors:** Lynn Buyachuihan, Yue Zhao, Christian Schelhas, Martin Grininger

## Abstract

Modular polyketide synthases (PKSs) are attractive targets for the directed, biosynthetic production of platform chemicals and pharmaceuticals by protein engineering. In this study, we analyze docking domains from the 6-deoxyerythronolide B synthase, SYNZIP domains, and the SpyCatcher:SpyTag complex as engineering tools to couple the polypeptides VemG and VemH to functional venemycin synthases. Our data show that the high-affinity interaction or covalent connection of modules, enabled by SYNZIP domains and the SpyCatcher:SpyTag complex, can be advantageous, e.g., in synthesis at low protein concentrations, but their rigidity and steric demand decrease synthesis rates. However, we also show that efficiency can be recovered when inserting a hinge region distant from the rigid interface. This study demonstrates that engineering approaches should take the conformational properties of modular PKSs into account and establishes a three-polypeptide split-venemycin synthase as an exquisite *in vitro* platform for the analysis and engineering of modular PKSs.

## INTRODUCTION

Polyketides are a structurally diverse class of natural products that comprise various functions in therapeutic and clinical applications, such as erythromycin, which is applied in antibiotic therapy, or rapamycin, which has immunosuppressant properties and is, for example, used to prevent organ rejection^1–4^. The complex scaffolds of polyketides are built from simple acyl building blocks by polyketide synthases (PKSs). PKSs can be divided into different types according to their mode of action and architecture. Type I modular PKSs belong to nature’s largest and most complex enzymatic factories^5^. They are hierarchically organized in modules and domains, whereby each module obligatory elongates the growing polyketide intermediate by two carbons and optionally further processes it upon passing it to the next module.

Modular PKSs work in an assembly-line like fashion and can be dissected into modules, each responsible for one round of chain elongation and processing. The growing intermediate is channeled from one module to the next until the end of the assembly line is reached, where the mature polyketide chain is released. A module includes at least three domains: the ketoacyl synthase (KS) domain, which is responsible for elongating the intermediate, the acyltransferase (AT) domain (acting in *cis* or *trans*), which selects the extender substrate (usually a malonyl- or methylmalonyl-CoA^5^), and an acyl carrier protein (ACP) which shuttles the intermediate between the catalytically active domains within and across modules^6^. The modules may harbor additional processing domains to set the reductive state and stereochemistry of the α- and β-position of the growing polyketide chain.

Modular PKSs are an exciting target for bioengineering due to the relatively strict correlation of protein organization and product structure. Known as the principle of collinearity ^2,7–14^, each chemical feature of a polyketide is directly linked to a specific module and domain(s). In this light, custom synthesis of polyketides with altered or improved functions seems possible through directed modifications of modular PKSs.

The venemycin PKS (VEMS) proved to be a suitable platform for investigating PKS engineering approaches (Figure 1A). Its small size facilitates its handling, the starter substrate and the final product absorb UV light, enabling direct activity monitoring by starter substrate depletion, and its turnover rate is higher than that of comparable simple modular PKS testbeds, allowing a more sensitive readout of the effects of engineering attempts on PKS activity^15,16^. VEMS was discovered in 2016 by activating a silent biosynthetic gene cluster of *Streptomyces venezuelae^17^*. The short modular assembly line is encoded by two polypeptides, VemG and VemH (Figure 1A). VemG comprises a loading module and an elongation module, and VemH a second elongation module and a C-terminal thioesterase domain (TE). The loading module starts with a PKS-untypical adenylation domain (A), which activates and transfers the 3,5-dihydroxybenzoic acid (DHBA) starter substrate onto the ACP. It also harbors an inactive KR domain that is thought to play a structural role^15,17^. Subsequently, the ACP-bound intermediate is passed onto the KS of the first elongation module, a step termed chain translocation. The KS domain catalyzes the decarboxylative Claisen condensation of the KS-bound intermediate with malonyl-ACP, that has been generated before by the malonyl-CoA-specific AT domain. The elongated intermediate is translocated to the second elongation module, where the next chain elongation step happens. As both elongation modules do not harbor processing domains, a triketide intermediate, tethered to the VemH ACP, is released by the TE domain upon cyclization to a pyrone ring, creating the product venemycin.

**Figure 1.**
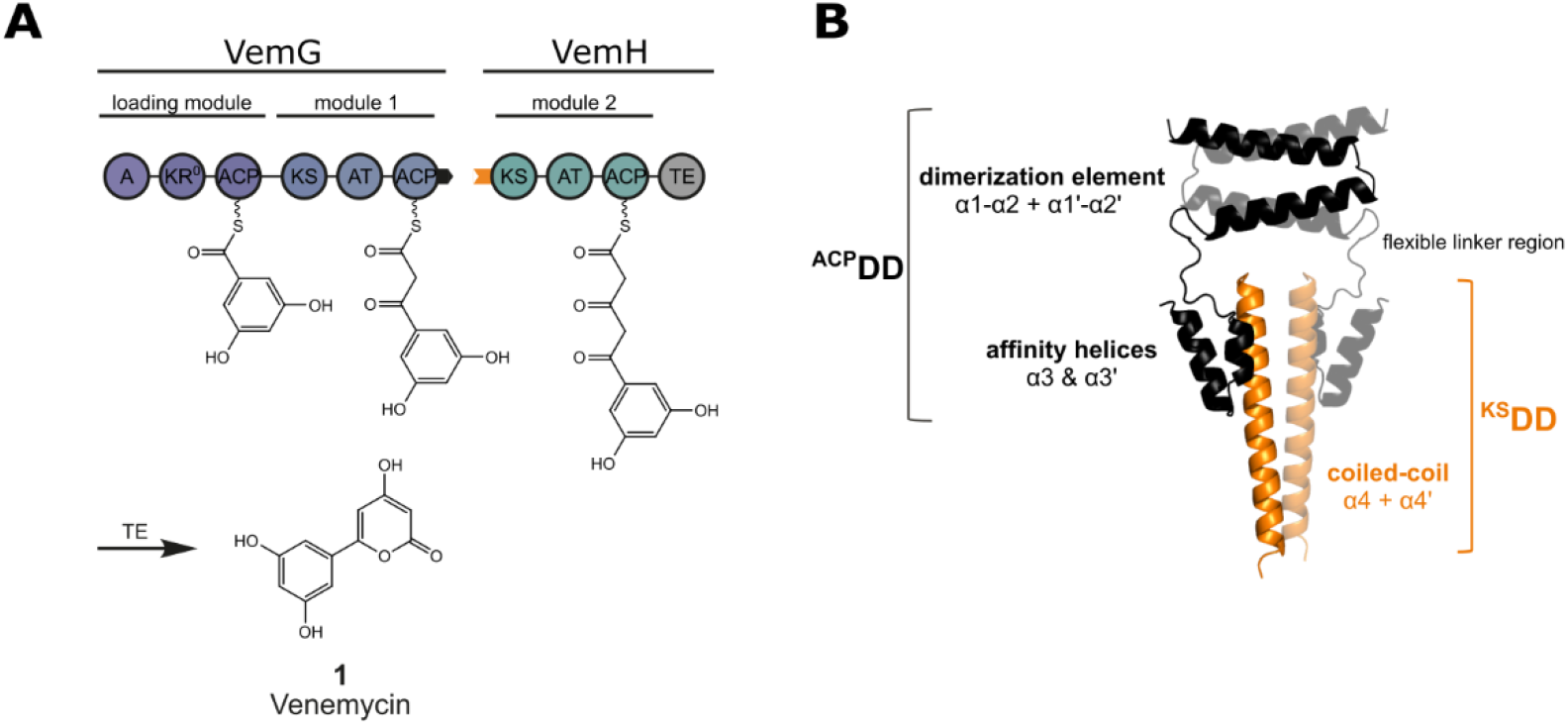
Module and domain organization of VEMS and modeled structure of the DDs mediating subunit communication between VemG and VemH. (A) The architecture of VEMS. The two polypeptides (VemG and VemH), the encoded elongation modules, as well as the loading module, the thioesterase domain (TE), and the final product venemycin (**1**) are depicted. Polyketide intermediates are attached to the respective acyl carrier protein (ACP). Black and orange tabs depict the ^ACP^DD and ^KS^DD, respectively. Domain annotations are as follows: A- adenylation domain, AT- acyltransferase, KS-ketoacyl synthase, ACP-acyl carrier protein, KR^0^- nonfunctional ketoacyl reductase, and TE-thioesterase. (B) Structural features of class 1 *cis*-AT PKS DDs. Prediction of the VemG:VemH docking interface using ColabFold^19^. The C-terminal ^ACP^DD comprises three α-helices and is depicted in black. While α1-α2 build a homodimeric dimerization element, the α3 affinity helix interacts with the ^KS^DD. The dimerization element and the affinity helix are connected by a 23-residue flexible linker region. ColabFold predicted a seven-residue helical region within the flexible linker region connecting the dimerization element and the affinity helix, which is no conserved feature of class 1 *cis*-AT PKS DDs and was therefore not considered in numbering the α-helices comprising the docking interface. The N-terminal ^KS^DD consists of one α-helix (α4), building a homodimeric coiled-coil, and is depicted in orange.

Successful chain transfer between VemG and VemH is only possible when the ACP of module 1 and the KS of module 2 are in spatial proximity, which is enabled by their C- and N-terminal docking domains (^ACP^DD and ^KS^DD, respectively, Figure 1B). The DDs of VEMS belong to the class 1 *cis-AT* PKS DDs, which are known to form a four α-helix bundle^18^. The ^ACP^DD comprises three α-helices, of which the first two helices form a homodimeric dimerization element, and the third helix is involved in non-covalent specific interactions with the complementary ^KS^DD. As PKS modules form homodimers, the N-terminal region is built by a total of six α-helices. The ^KS^DD consists of a single α-helix and forms a homodimeric coiled-coil structure. Current data suggest that PKS DDs interact with a K_D_ in the low μM range^18^; e.g., the K_D_ of the venemycin PKS subunits was determined to be 5.1 μM^15^. The high dissociation constants, with which protein subunits are assembled to modular PKSs, inspired researchers to test different docking tools with higher affinity to increase the efficacy of modular PKSs and facilitate *in vitro* handling. Higher affinity docking tools enable running the assembly line at lower enzyme concentrations by increasing the portion of assembled protein subunits.

In this study, we investigated docking tools for their ability to replace the native DDs of modular PKSs. The tested tools varied in affinity, structure, and size. We chose VEMS as a testbed because it possesses just one native DD-mediated chain translocation interface between VemG and VemH. Overall, we replaced the native DDs with three different types of interaction domains, each of them well characterized properties: (i) the homologous class 1 *cis*-AT PKS DDs from DEBS^20,21^, (ii) the meanwhile in megasynthase engineering well-established SYNZIP domains ^16,22,23^, as well as (iii) the SpyCatcher:SpyTag complex^24^.

All tested DDs introduced a bottleneck at the VemG:VemH interface, emphasizing that the structural features of the venemycin DDs well match the kinetic requirement of the translocation reaction. Replaced by SYNZIPSs, a kinetic penalty arose, most likely from the long and rigid coiled-coil structural element that restricts the conformational variability of the modPKS in general and the interplay of domains during chain translocation in particular. Similarly to the SYNZIPSs, also the SpyCatcher:SpyTag complex decreased turnover rates, which may again arise from steric constraints. Nevertheless, our study reveals the SpyCatcher:SpyTag complex as a valuable tool in the engineering of PKSs and presumably also of the related non-ribosomal peptide synthases (NRPSs). Based on the ability to glue proteins by isopeptide bond formation, the covalently linked, single-polypeptide VEMS ran at turnover rates independent of protein concentrations. Thus, the SpyCatcher:SpyTag complex offers the chance to increase product output when operating assembly lines at low subunit concentrations.

Finally, we created a three-polypeptide VEMS (split VEMS) by introducing a DEBS-derived docking interface into the VemG subunit between the loading module and module 1. The smaller polypeptides showed improved properties in recombinant production, while the split VEMS exhibited the same turnover rate as the native PKS *in vitro*. Intriguingly, the DEBS-derived docking interface within VemG (between the loading module and module 1) relieved the kinetic penalty at the VemG:VemH translocation interface introduced by the SYNZIPSs, presumably by releasing the conformational constraints of the zipped interface. With AlphaFold2-based modeling, using ColabFold^19^, we visualized the structural requirements of the translocation reaction. Restricted by interdomain linkers, the interaction of the ACP with the KS domain during polyketide translocation seems just possible when larger conformational changes of the modules accompany the ACP mobility.

Overall, our findings establish VEMS as a suitable *in vitro* platform for studying modular PKSs, and enlarges the toolbox of interaction domains by the SpyCatcher:SpyTag complex. In demonstrating that PKS engineering should take overall conformational properties into account, our study further highlights a yet underexplored aspect of PKS engineering.

## RESULTS AND DISCUSSION

### Testing Communication Tools at the VemG:VemH Translocation Interface-

The subunits VemG and VemH were produced in *Escherichia coli*. The two proteins forming the VEMS assembly line were used at a final concentration of 8 μM, above the reported K_D_ value of 5.1 μM^15^. The activity of the native PKS was analyzed *in vitro* by an HPLC-based assay, monitoring the starter substrate consumption via its UV absorbance. Determination of the PKS activity at different temperatures revealed a temperature dependence of the rate-limiting step of the venemycin synthesis. The highest activity of 6.3 min^-1^ was observed at 37 °C (Figure S1). With the *in vitro* assembly line in hand, we first sought to assess the importance of the DDs for synthesis. Thus, we removed the DDs, that naturally mediate communication at the VemG:VemH interface, to receive a DD-less VEMS. Intriguingly, deleting the ^ACP^DD, deprived VemG of the ability to adopt an active dimeric state required for PK synthesis (Figure S2) and resulted in a dramatic loss of activity of the DD-less VEMS. The DD-less VEMS was able to produce venemycin, as confirmed by LC-MS analysis of the reaction mixture (Figure S3), but at just 1% of the activity of the native VEMS (2, Figure 2B). Similar effects were previously observed for other PKSs^25^, underscoring the importance of DDs in colocalizing protein subunits to increase the effective concentration of domain interaction partners within and across modules. We investigated three different interaction domains in their ability to replace the VEMS DDs. Among them DEBS-derived DDs, SYNZIPSs, and the SpyCatcher:SpyTag complex, each of them well understood in their properties. The interaction domains cover a wide spectrum of thermodynamic (K_D_) and structural properties (Figure 2A). In a first analysis of the VemG:VemH subunit interaction, we replaced the native DDs with the homologous DD pair from the 6-deoxyerythronolide B synthase (DEBS) that natively mediates communication between the proteins DEBS1 and DEBS2. The DEBS-derived DDs belong to the same class of DDs (class 1 *cis*-AT PKS DDs), which are known to possess conserved structural features and interact with comparable affinity within the low μM range^15,18,20^.

**Figure 2.**
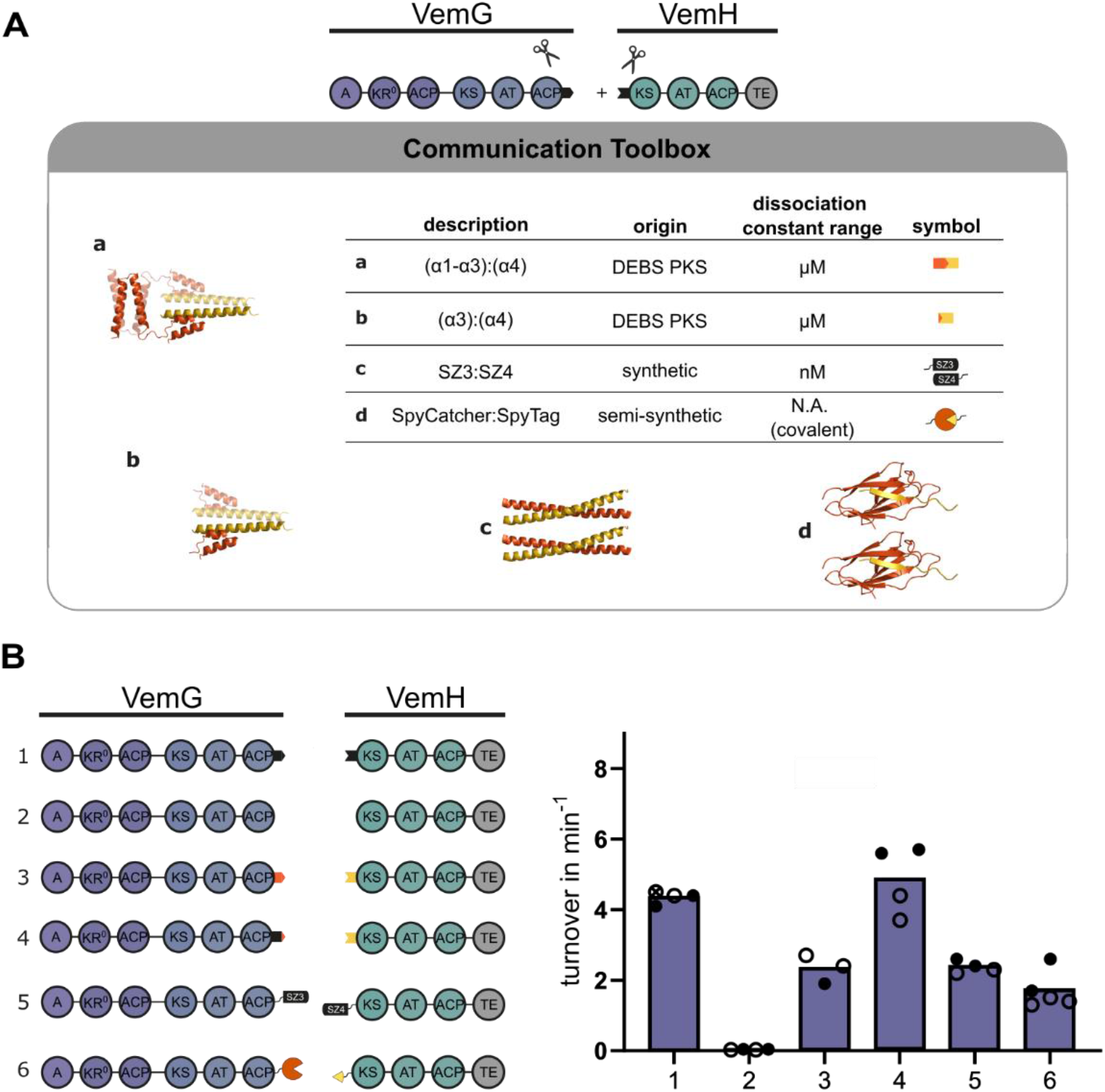
Communication tools for engineering VemG:VemH subunit communication. (A) Design of the docking study. Docking tools fused to VemG and VemH are depicted in orange and yellow, respectively. Two different designs for the ^ACP^DD swap, one replacing the whole DD (a) and one replacing just the affinity helix (b). The structural DD parts which were replaced are shown by means of the VemG:VemH docking interface prediction with ColabFold^19^. c: The high affinity SYNZIP pair SZ3 and SZ4. Shown is the structure of the SZ1:SZ2 pair (PDB ID: 3HE5)^29^, as there is no structure available for the SZ3:SZ4 pair. The SZ1:SZ2 pair forms a dimeric coiled-coil structure in a parallel orientation as the SZ3:SZ4 pair. d: The SpyCatcher:SpyTag complex (PDB ID: 4MLI)^30^ which forms by covalent connection within minutes. (B) The native VEMS and five chimeric PKS bearing an engineered VemG:VemH translocation interface. Within the text the PKSs are named as follows: 1 – VEMS, 2 – DD-less VEMS, 3 – (α1-α3)-swapped VEMS, 4 – (α3)-swapped VEMS, 5 – SYNZIP-linked VEMS, and 6 – Spy-linked VEMS. Turnover rates were determined at an enzyme concentration of 8 μM at 30 °C. Biological replicates are shown as dots of different color. Removing the docking interface (2) diminished the activity of VEMS. While the DEBS α3-swap (4) resulted in a comparable activity to the native VEMS, the synthetic docking tools exhibited around half of the activity of the native VEMS.

In VemH, we replaced the N-terminal α-helix, which is termed α4 helix or ^KS^DD. It interacts with the C-terminal α-helix of VemG, which is the affinity helix (α3) of docking domain termed ^ACP^DD. The ^ACP^DDs of this DD class comprises an additional dimerization element α1-α2 N-terminal to α3 (Figure 1B); thus, we aimed for two designs for the ^ACP^DD-swap (Figure 2A). In the first design, we exchanged the entire ^ACP^DD (termed (α1-α3)-swapped VEMS), including the dimerization element (α1-α2) and the affinity helix (α3). In the second design ((α3)-swapped VEMS), we solely replaced the affinity helix, assuming that the ^ACP^DD dimerization element is not involved in ^ACP^DD:^KS^DD interactions^18^. While the (α1-α3)-swapped VEMS exhibited about half of the activity of the native VEMS (3, Figure 2B), the (α3)-swapped VEMS was slightly faster than the native assembly line (4, Figure 2B).

Next, we used the synthetic SYNZIPs^22^ (SZ3:SZ4, Figure 2A) for docking, which were well-established during the recent years as tools in modular PKSs^26,27^ and NRPSs^23,28^ engineering. The selected SYNZIP pair SZ3:SZ4 forms a parallel dimeric coiled-coil structure with a K_D_ below 30 nM^22^. Bridging the VemG:VemH translocation interface with the SZ3:SZ4 pair resulted in 55% activity of the native PKS (5, Figure 2B). In the native docking interface, a flexible linker region connects the dimerization and the affinity element (Figure 1B, 23 residues for VEMS according to ColabFold^19^ prediction), allowing flexible movement of the linked modules to each other. In contrast, the artificial VemG:VemH translocation interface (Figure S4), generated with the approx. 60 Å coiled-coil forming SYNZIPSs, is rather rigid, which may hamper polyketide translocation between VemG and VemH and reduce turnover rates.

As a third tool, we tested a covalent bond-forming tag system (SpyCatcher:SpyTag complex, Figure 2A) to enable communication between the protein subunits VemG and VemH. The SpyTag:SpyCatcher complex is an engineered bacterial adhesin, consisting of a 13-residue peptide tag (SpyTag) and a 15 kDa protein partner (SpyCatcher), that enables irreversible linkage between the tagged proteins within minutes by spontaneous isopeptide bond formation between a Lys of the SpyCatcher and an Asp of the SpyTag^24^. To the best of our knowledge, the SpyCatcher:SpyTag complex has not yet been used for megasynthase engineering, nor has it been applied for other polypeptides of this size. We fused the SpyCatcher to the C-terminus of VemG downstream of the ^ACP^DD dimerization element (α1-α2, Figure 1B) to preserve the dimerization-promoting effect of the dimerization element on the VemG subunit. The Spy-tagged VemH was generated by replacing the ^KS^DD (α4, Figure 1B) with the SpyTag sequence (Figure 2B). The irreversible covalent linkage of the protein subunits was tracked by SDS PAGE (Figure 3A), revealing that the covalent bond formation was completed within 10 minutes. The covalently linked assembly line exhibited 39% of the activity of the native PKS (6, Figure 2B).

**Figure 3.**
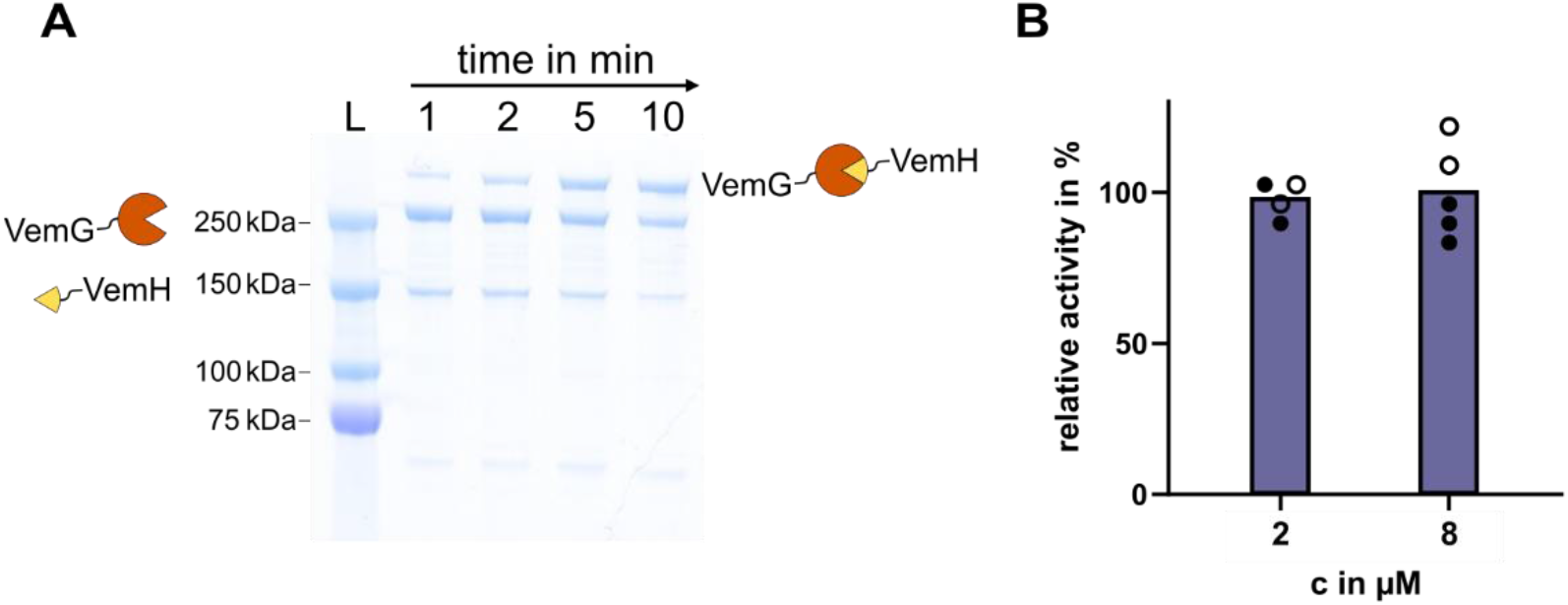
Characterization of the Spy-linked VEMS. (A) SDS-PAGE of the SpyCatcher:SpyTag reaction. The 15 kDa SpyCatcher is indicated in orange at the C-terminus of VemG (246 kDa), while the SpyTag is indicated in yellow at the N-terminus of VemH (141 kDa). Within one minute, a band representing the Spy-linked VEMS (387 kDa) appeared (lane 1). Under the given assay conditions, the reaction is completed within 10 minutes. (B) Relative activity of the Spy-linked VEMS at different enzyme concentrations and 30 °C reaction temperature. The Spy-linked VEMS can be operated at low enzyme concentrations without loss in activity as PKS subunit dissociation is prevented by covalent linkage.

To better understand how the docking tools impact the chain translocation reaction, we analyzed the native VemG:VemH docking interface in its structural requirements and steric constraints. A multiple sequence alignment of a set of ACP-DD sequences of class 1 *cis*-AT PKSs revealed that the linker connecting the ACP and the ^ACP^DD is typically around 8-13 residues long (Figure S5), which is relatively short, considering that the linker must provide sufficient conformational freedom to the ACP to shuttle the polyketide across the module:module interface. In this regard, the physical interaction of the ACP and the KS domain of the downstream module does not seem possible by the positional variability of the ACP alone but requires the flexibility of adjacent domains to drag the ACP across the interface. We propose that the linker region, connecting the ^ACP^DD dimerization element and the affinity helix (Figure 1B) acts as flexible waist that is crucial for the overall flexibility of VEMS and modular PKSs in general^31^. For visualization of the structural requirements of the chain translocation reaction, we predicted the DEBS ACP4-DD:DD-KS5 translocation interface with ColabFold^19^ (Figure S6). The choice fell on this interface as except for the ACP4 the structures of all other domains comprising this interface are already solved (PDB(docking domains): 1PZQ+1PZR, PDB(KS5): 2HG4)^32,33^. Four of the five best-ranked structures predicted the ACP4 docked to the KS5 active site entry. All of these four structures seem plausible given that the serine of ACP, that harbors the phosphopantetheine arm for substrates shuttling, is positioned in suitable distance to the KS active site. The models indicate that the ACP is indeed unable to reach the KS active site without collaborative conformational dynamics of the involved modules.

### The One-Polypeptide Assembly Line-

Although the native VEMS outperforms the Spy-linked VEMS under the given assay conditions (c(enzyme) = 8 μM, Figure2C), the covalent connection can be harnessed to maintain the activity of the assembly line at lower enzyme concentrations (below K_D_). While dilution of the native VEMS leads to a dramatic drop in activity due to dissociation of the assembly line-forming subunits (Figure S7)^15^, the Spy-linked VEMS can be operated at lower concentrations without loss of activity (Figure 3B), as the irreversible covalent linkage ensures a high effective concentration of the domains interacting during chain transfer at the VemG:VemH interface. Particularly in the field of megasynthase engineering, the SpyCatcher:SpyTag complex is an elegant tool that allows the fusion of protein subunits post-translationally. This approach can likely be transferred to other members of the megasynthase family, like NRPSs, where subunit interaction is also mediated by DDs that interacting with weak affinity^18,34,35^.

### PKS Splitting Improves Protein Yield and Purity Without Harming Activity-

The production of large proteins in heterologous hosts like *E. coli* is often accompanied by low yields, proteolytic degradation, and poor protein quality^20,36^. We thought that the access to VEMS and its suitability as an *in vitro* testbed for analyzing modular PKSs could be improved by splitting the first polypeptide VemG (236 kDa) into its loading module and the first elongation module and linking them non-covalently by DDs. We introduced the class 1 *cis-AT* DDs from DEBS, as they naturally occur at ACP:KS translocation interfaces of actinobacterial modular PKSs^18^ (Figure 4A). This approach facilitates the VEMS purification process and enables harnessing VEMS in a mix-and-match approach with already available PKS testbeds, as all three modules comprising VEMS exist separately. We chose DDs from the DEBS pathway, which natively bridge the translocation interface between DEBS2 and DEBS3 because they were previously applied to an engineered DEBS assembly line reconnecting the separated loading module and the first elongation module^20^. The native 31-residue linker connecting the loading module and module 1 of VemG was removed and the modules were non-covalently reconnected by introducing the DEBS-derived DDs which natively bridge the DEBS2:DEBS3 translocation interface. The seven amino acid long linker, natively connecting DEBS ACP4 and its C-terminal ^ACP^DD was included to ensure that the Vem ACP0 is sufficiently mobile to reach the Vem KS1 for polyketide translocation. The resulting two smaller subunits could be produced in *E. coli* at twofold and fourfold increased yields for the isolated loading module and module 1, respectively (Figure 4C). In activity assays, again monitoring the starter substrate consumption via HPLC, the split VEMS proved to be as fast as the native VEMS (Figure 4B). The data imply that the translocation across the newly installed non-covalent interface is not invasive or at least not introducing a kinetic bottleneck in the venemycin synthesis.

**Figure 4.**
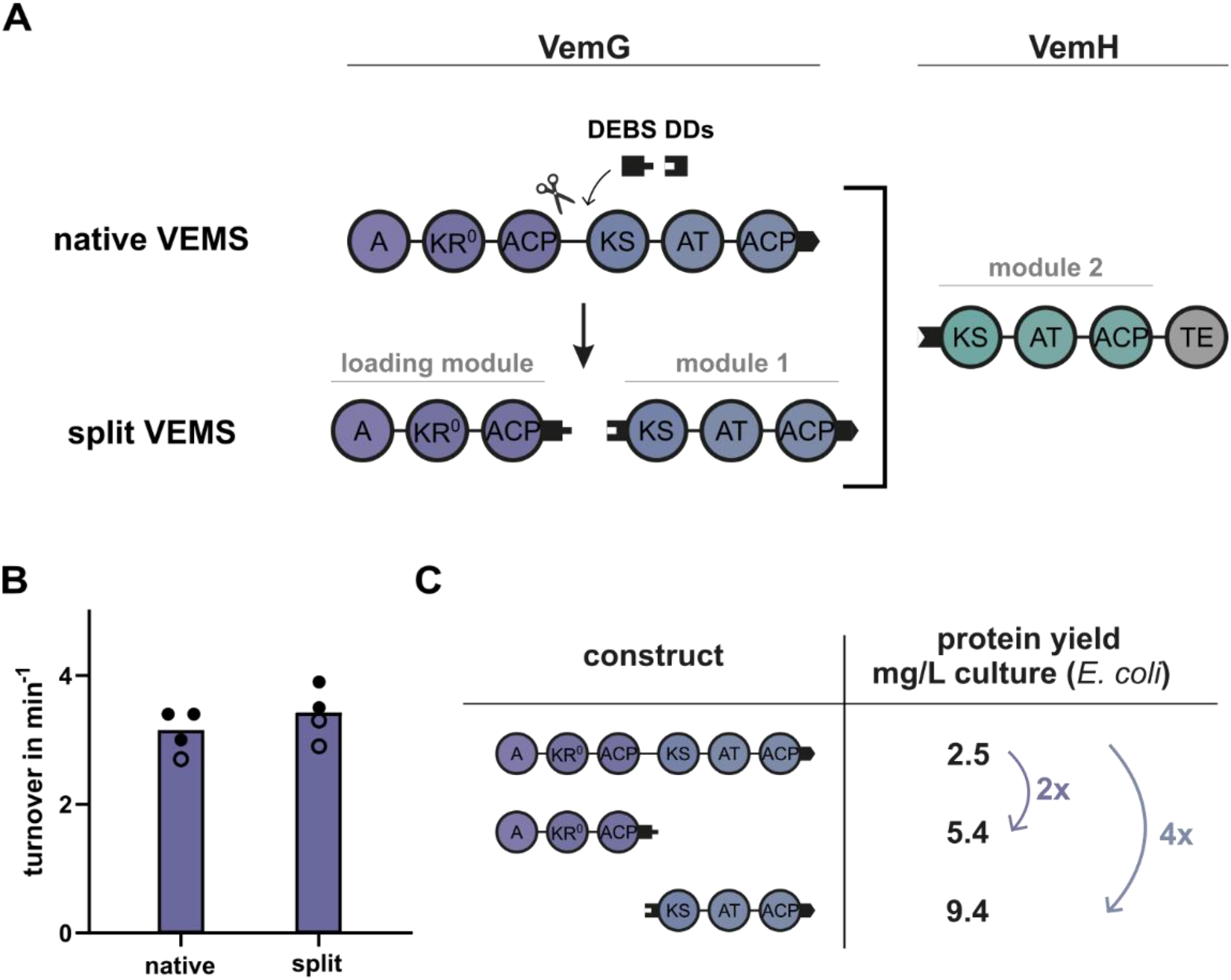
Module organization and characterization of the split VEMS. (A) VemG was separated into its loading module and the first elongation module at the translocation interface between the ACP of the loading module (ACP0) and the KS domain of the first elongation module (KS1). The PKS activity was regained by introducing DEBS-derived DDs (depicted as black tabs). (B) Turnover of the native and the split VEMS at 8 μM enzyme concentration and 25 °C. Introducing a non-native docking interface within VemG has nearly no influence on PKS activity. (C) Purification yields of the native VemG and the separate modules comprising VemG. Purification yields improved for the smaller proteins.

### Splitting Engineered PKS Can Improve Turnover-

Since the introduction of a non-native docking interface within VemG proved to be successful (split VEMS, Figure 4), we sought to investigate whether the SYNZIP-linked VEMS could benefit from splitting VemG. Given the rigidity of the SYNZIP domains, which we assumed to decrease translocation efficiency between VemG and VemH, we hypothesized that we could regain PKS activity by introducing conformational variability by introducing class 1 *cis*-AT PKS DDs upstream of the rigid SYNZIP interface.

To visualize the SYNZIP-mediated module:module interaction, we used ColabFold^34^ to predict the translocation interface built by the split SYNZIP-linked VEMS subunit parts ACP1-SZ3 and SZ4-KS2 (Figure S4). The structure model revealed that the ACP can just reach the KS active site entrance under extensive bending of the SYNZIP interface towards the KS2 dimer and excludes the possibility of a linear organization of the interacting modules during the chain translocation step. The structure prediction supports the hypothesis that the bottleneck introduced by the SYNZIP domains originates from their rigidity. The C-terminally SZ3-tagged VemG was split into its loading module and the elongation module, analogously to the split VEMS, to receive the split SYNZIP-linked VEMS (Figure 5A). Indeed, the new interface in the SYNZIP-linked VEMS increased the activity of the assembly line 3.5-fold (Figure 5B). The recovery of activity by engineering VEMS off-site, is an intriguing example for the complexity of the module:module and domain:domain interplay within modular PKSs.

**Figure 5.**
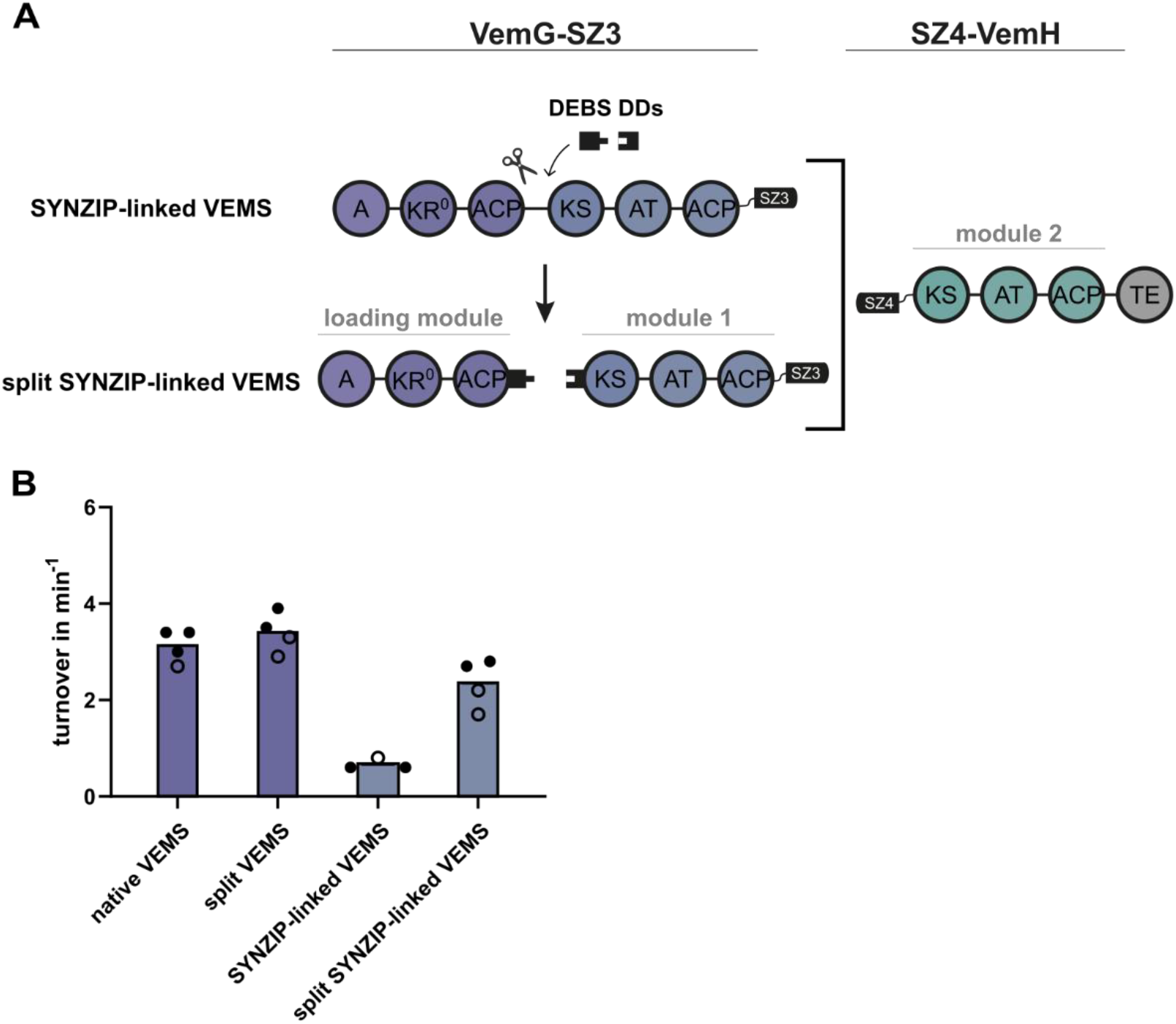
The schematic architecture and characterization of the split SYNZIP-linked VEMS. (A) VemG was separated into its loading module and the first elongation module at the translocation interface between the ACP of the loading module (ACP0) and the KS domain of the first elongation module (KS1). The communication between the modules was restored by introducing DEBS-derived docking domains (depicted as black tabs), which natively enable docking between the polypeptides DEBS2 and DEBS3. Docking at the VemG:VemH translocation interface is enabled by SYNZIPs. The C-terminal DD of VemG is replaced by SZ3, while SZ4 replaces the N-terminal DD of VemH. (B) Turnover of the native and the split VEMS and turnover of the SYNZIP-linked and split SYNZIP-linked VEMS at 8 μM enzyme concentration and 25 °C. Introducing a non-native docking interface within VemG has nearly no influence in the context of the native VEMS but can improve turnover in the SYNZIP-linked VEMS 3.5-fold.

## Conclusion

Modular PKSs are among the most intricate enzymes on earth. Their hierarchical organization in modules and domains promises that polyketides can be regioselectively modified by directed changes in the assembly line arrangement, eventually enabling the programmable production of designer molecules. Many years of progress in the structural elucidation of domains and subregions of PKS modules, the recently acquired knowledge about the structure of PKS modules^37–40^, and the new possibilities of accurate structure predictions (AlphaFold2^41^, RoseTTAFold^42^) currently cumulate in an advanced structural understanding of modular PKS-mediated assembly line synthesis. Progress in modular PKSs engineering now depends on quantitative, *in vitro* enzymatic studies to better understand the recent findings on the complex interplay of substrate specificity and domain:domain interactions and to develop broadly applicable engineering strategies ^15,20,27,43,44^. Unfortunately, purifying PKS polypeptides in high yields and high quality is difficult because the large size of the proteins (hundreds of kDa) hampers successful production in heterologous hosts^20,36^. For example, to reconstitute modular PKSs in sufficient amounts from recombinantly expressed subunits, polypeptides harboring two or more modules were split into shorter polypeptides ^45,20^. Overall, just a handful of modular PKSs are available *in vitro*, that are suitable for quantitative studies of which DEBS as well as truncated versions thereof, have been used most frequently^20^.

In this study, we worked with the recently discovered venemycin synthase (VEMS) ^17^ and further developed its use as a modular PKS testbed. VEMS is generally well-suited for *in vitro* studies because it is small, fast in turnover, and well-accessible by recombinant production^15^. With various engineered variants of VEMS, we set out to test interaction domains (naturally occurring in modular PKSs (class 1 *cis*-AT PKS DDs^18^) as well as synthetic domains used as engineering tools (SYNZIPs^22^ and SpyCatchter:SpyTag complex^24^) in their impact on assembly lines synthesis. While all tested tools enabled communication between the VEMS subunits, our data suggest that the inherent flexibility of class 1 *cis*-AT PKS DDs is vital for a fast chain translocation reaction. Available synthetic docking tools are superior in gluing modules either by high affinity or covalently but cannot provide sufficient flexibility, hampering a fast intermodular chain translocation. Our study also provides a possible solution to this problem. We demonstrate that the bottleneck introduced by the rigid SYNZIPs can be released when adding new flexibility to the assembly line. Specifically, the SYNZIP-linked VEMS recovered high turnover rates when introducing a hinge region within VemG by the conformationally flexible PKS DDs. Notably, recent structural studies reported the high conformational variability of a PKS module^37–40^, which extends to the whole assembly line^46^ to enable the translocation of the polyketide intermediate. Thus, our data on the importance of the relative conformational variability of PKS modules align well with the current structural understanding of modular PKSs.

In sum, our study on the comparison of docking domains suggests that engineering strategies in modular PKSs should keep conformational dynamics as an important parament in mind. Moreover, our data encourage future developments of communication tools for modular PKSs that take structural flexibility into account. Finally, the transformation of the native two-polypeptide VEMS into the tri-polypeptide split VEMS not only supported our understanding of then conformational properties of modular PKSs but also generated a new version of the venemycin synthase that is interesting for the chemical biological community. The split VEMS is as fast as the wild-type version but accessible with higher yields and a more versatile PKS analysis and engineering platform.

## METHODS

### Reagents

CloneAmp HiFi PCR Premix was from Takara. Primers were synthesized by Sigma Aldrich. For DNA purification, the GeneJET Plasmid Miniprep Kit and the GeneJET Gel Extraction Kit from ThermoFisher Scientific were used. Cloning was performed with the In-Fusion HD Cloning Kit from Takara. NiCo21 (DE3) Competent cells were from New England BioLabs. Stellar Competent Cells were from Takara and One Shot BL21 (DE3) Cells were from ThermoFisher Scientific .All chemicals for buffer preparations were from Sigma-Aldrich. LB (Lennox) and 2 xYT media for cell cultures were from Carl Roth. Isopropyl-β-D-1-thiogalactopyranoside (IPTG), ampicillin, and carbenicillin were from Carl Roth. Spectinomycin was from Sigma-Aldrich. His60 Ni Superflow Resin was from Takara, and Strep-Tactin Sepharose columns were from IBA Lifesciences. For anion exchange chromatography, the HiTrapQ HP column (column volume = 5 mL) was from Cytiva. Proteins were concentrated with Amicon Ultra centrifugal filters from Merck Millipore. Coenzyme A and 3,5-dihydroxybenzoic acid were from Sigma-Aldrich. Malonic acid and magnesium chloride hexahydrate were from Carl Roth. Adenosine-5’-triphosphate (ATP) was from Sigma-Aldrich. Reducing agent tris-(2-carboxyethyl) phosphine (TCEP) was from ThermoFisher Scientific.

### Plasmids

The DNA encoding VemG and VemH was amplified from the genomic DNA of *Streptomyces venezuelae* ATCC 10712 (DSMZ) by PCR and introduced into a pET22b(+) expression vector by In-Fusion Cloning (Takara). These expression plasmids were used as a template to generate all engineered venemycin assembly line constructs of this study via In-Fusion Cloning (Takara). The resulting plasmids, cloning strategies, and primer sequences are further specified in Tables S1, S2 & S3. The plasmid sequences were verified by Sanger Sequencing (Microsynth Seqlab). All protein sequences of the constructs generated in this study are listed in Tables S4 & S5.

### Bacterial Cell Cultures and Protein Purification

All PKS proteins were expressed and purified using similar protocols. For protein expression, different *E. coli strains* were tested, revealing that the expression strain had no influence on the activity but on the yield of the proteins. Finally, VemG-based constructs were expressed in BAP1^47^, VemH-based constructs and (5)M1 in NiCo21, and LM(4) in BL21 cells. All proteins were expressed in the *holo-form* (to activate the ACP domain post-translationally with a phosphopantetheine arm). For expression in NiCo21 and BL21 cells, the construct-encoding plasmids were co-transformed with a plasmid encoding for the phosphopantetheine transferase Sfp from *B. subtilis* (pAR357ref). Cell cultures were grown on a 2 L scale in 2x YT media at 37 °C until an OD_600_ of 0.3 was reached, whereupon the temperature was adjusted to 18 °C. At an OD600 of 0.6, protein production was induced by adding 0.1 mM IPTG, and the cells were grown for another 18 h. Cells were harvested by centrifugation at 5,000 xg for 15 min and lysed by French Press (lysis buffer: 50 mM sodium phosphate, 10 mM imidazole, 450 mM NaCl, 10% glycerol, pH 7.6). The cell debris was removed by centrifugation at 50,000 xg for 45 min. The supernatant was purified using affinity chromatography. All constructs contained a C-terminal His-tag, and VemG-based constructs contained an additional N-terminal twinstrep-tag for tandem purification.

For proteins containing just a C-terminal His-tag, the purification procedure was as follows: The supernatant was applied onto the column (5 mL Ni resin). A first wash step was performed with the above-mentioned lysis buffer (10 column volumes), followed by a second wash step with 10 column volumes of wash buffer (50 mM sodium phosphate, 25 mM imidazole, 300 mM NaCl, 10 % glycerol, pH 7.6). Proteins were eluted with 6 column volumes of elution buffer (50 mM sodium phosphate, 300 mM imidazole, 10% glycerol, pH 7.6). The eluate was purified by anion exchange chromatography using a HitrapQ column on an ÄKTA FPLC system (column volume 5 mL). Buffer A consisted of 50 mM sodium phosphate, 10% glycerol, pH 7.6, whereas buffer B contained 50 mM sodium phosphate, 500 mM NaCl, 10% glycerol, pH 7.6.

For proteins containing a C-terminal His- and an N-terminal twinstrep-tag, the purification procedure was as follows: The supernatant was applied onto the first affinity chromatography column (5 mL Ni resin). One wash step was performed with 5CV of the above-mentioned lysis. Proteins were eluted with 2×2.5 column volumes of elution buffer (50 mM sodium phosphate, 300 mM imidazole, 10% glycerol, pH 7.6). The eluate was applied to the second affinity chromatography column (5mL strep resin) and washed with 6 CV strep-Wash buffer (50 mM sodium phosphate, 10% glycerol, pH 7.6). The target protein was eluted with 2×2.5 CV strep-elution buffer (50 mM sodium phosphate, 2.5 mM desthiobiotin, 10% glycerol, pH 7.6). The eluate was purified by anion exchange chromatography using a HitrapQ column on an ÄKTA FPLC system (column volume 5 mL). Buffer A consisted of 50 mM sodium phosphate, 10% glycerol, pH 7.6, whereas buffer B contained 50 mM sodium phosphate, 500 mM NaCl, 10% glycerol, pH 7.6. The enzyme MatB (extender substrate regeneration system) was purified as described previously^20^. Protein concentrations were determined with a Nanodrop. Samples were stored as aliquots at −80 °C until further use.

### Size Exclusion Chromatography

To determine the purity and the oligomeric state of the proteins after ion exchange chromatography, samples of each protein were analyzed by size exclusion chromatography on an ÄKTA FPLC system using a Superose 6 Increase 10/300 GL column from Cytiva (buffer: 50 mM sodium phosphate, 500 mM NaCl, 10% glycerol, pH 7.55). The SEC profiles and SDS PAGE images of all proteins are provided in Figures S2 & S8.

### Thermal Shift Assay

The melting temperatures of all proteins were determined via Thermofluor assay^48^. The proteins were measured with a final concentration of 1.2 mg/mL in a buffer consisting of 50 mM sodium phosphate, 500 mM NaCl, and 10% glycerol at pH 7.6. Measurements were performed in technical duplicates. The melting temperatures are provided in Table S6.

### HPLC Activity Assay

All assays were performed in the reaction buffer (400 mM sodium phosphate, 10 % glycerol and pH 7.2) at different temperatures (the reaction temperature of each assay is reported in the corresponding figure caption). The reducing agent TCEP was used in a final concentration of 5mM. *In situ* extender substrate generation was performed using 10 μM MatB, 10 mM malonate, and 1 mM CoA. ATP and MgCl2 were provided in a final concentration of 9 mM. The starter substrate DHBA was used in a final concentration of 0.75 mM. If not otherwise stated, the enzymes constituting the assembly line were used in a final concentration of 8 μM. The reaction and protein solutions were incubated separately at the reaction temperature for 5 minutes. The reaction was started by pooling both solutions. 25 μL of the reaction solution was quenched by adding 5 μL of 70% perchloric acid and neutralized with 5 μL 10 M NaOH at six different time points within 10-20 minutes. After centrifugation at 20,000 xg for 5 minutes at 4 °C, 10-15 μL of the supernatant was injected into the HPLC. HPLC measurements were performed with a C18 Syncronis column from ThermoFisher Scientific from 5-60% MeOH in ammonium acetate (200 mM pH 6.0). The starter substrate consumption was tracked at 300 nm, and extender substrate regeneration was monitored at 260 nm. Within the study, the assay has proven to be reliable with only minor deviations between activities determined for biological duplicates with one exception. For one biological replicate of the (α1-α3)-swapped VEMS an exceptionally high activity (nearly threefold as active as the residual replicates) was determined which was excluded in Figure 2 and the grand mean calculation of the activity of the (α1-α3)-swapped VEMS.

### Liquid Chromatography-Mass Spectrometry Analysis of the HPLC assay reaction mixture

For LC-MS analysis, 40 μL of the HPLC assay reaction mixture was incubated overnight at room temperature and extracted two times with 450 μL ethyl acetate. Dried samples were reconstituted in 100 μL methanol and analyzed by HPLC-MS or HPLC-HRMS in positive or negative mode. Components were separated over a 16 min linear gradient of acetonitrile from 5% to 95% in water. Venemycin could be detected in positive and negative mode. *m/z* ratio of **1** [M+H]^+^ m/z = 221.05 and [M-H]-m/z = 219.03).

### Bioinformatical Analysis

Sequence alignments were generated with *Jalview* 2.10.5 (ClustalWS, default settings)^49^. Structure predictions were performed with ColabFold^19^ using default settings. The complexes were predicted without template information. MSA options were set as follows: MMseqs2 (UniRef+Environmental) was chosen as MSA mode and unpaired+paired as pair mode, and the model type was set to auto. Amino acid sequences used for complex prediction are provided in Tables S7, S8 & S9. AlphaFold error estimates are provided in Figures S9, S10 & S11.

## Acknowledgments

We thank Helge Bode for use of their LC-MS instruments and Jacob Hirschberg for support during his master thesis.

## Authors contribution

L.B. and M.G. wrote the manuscript. L.B. conceived and supervised the project and designed the constructs. L.B., Y.Z., and C.S. cloned and purified the constructs, analyzed the activity, thermal stability, and oligomerization. M.G. designed the research and analyzed the data.

### Supporting Information Available

Sequence alignment, structural models, additional activity analysis, SEC data, LC-MS data, list of plasmids, construct design, protein sequences and construct melting temperatures. This material is available free of charge via the internet at http://pubs.acs.org.

