## Supplementary Information for "Docking Domain Engineering in a Modular Polyketide Synthase and its Impact on Structure and Function"

### Content

**Supplementary Figures:**

Figure S1. Turnover rates of the native VEMS at different temperatures.
Figure S2. Analysis of proteins by SEC – VemG and VemG-based constructs.
Figure S3. LC-MS analysis of the venemycin production.
Figure S4. VemG-SZ3:SZ4-VemH interface predicted with ColabFold.
Figure S5. Alignment of ACP-ACPDD sequences from class 1 cis-AT PKS.
Figure S6. DEBS2:DEBS3 translocation interface.
Figure S7. Activity of the native VEMS at different enzyme concentrations.
Figure S8. Analysis of proteins by SEC – VemH and VemH-based constructs.
Figure S9. Quality of the native VemG:VemH docking domain interface predicted with ColabFold.
Figure S10. Quality of the DEBS2:DEBS3 docking domain interface predicted with ColabFold.
Figure S11. Quality of the VemG-SZ3:SZ4-VemH interface predicted with ColabFold.

**Supplementary Tables:**

Table S1. Cloning strategy of plasmids generated in this study.
Table S2. DNA and amino acid sequences of plasmids which are not published elsewhere and were used as a template to generate the plasmids of the constructs used in this study
Table S3. Plasmids used in this study and their origin
Table S4. Amino acid sequences of VemG and VemG-based constructs
Table S5. Amino acid sequences of VemH and VemH-based constructs
Table S6. Melting temperatures of the proteins composing the native and engineered VEMSs determined by thermal shift assay
Table S7. Amino acid sequences used for the complex prediction of the native VemG:VemH docking interface with ColabFold
Table S8. Amino acid sequences used for the complex prediction of the SYNZIP VemG-SZ3:SZ4-VemH docking interface with ColabFold
Table S9. Amino acid sequences used for the complex prediction of the DEBS2:DEBS3 docking interface with ColabFold

### Supplementary Figures

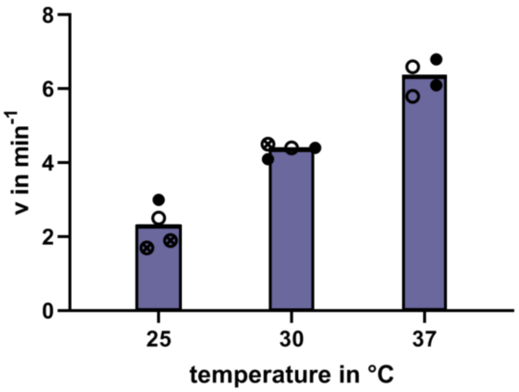

**Figure S1. Turnover rates of the native VEMS at different temperatures.** Data were obtained at a PKS concentration of 8 µM. Dots of different fillings represent biological replicates (independently purified protein samples). The rate-limiting reaction in the multistep PK synthesis depends on the reaction temperature.

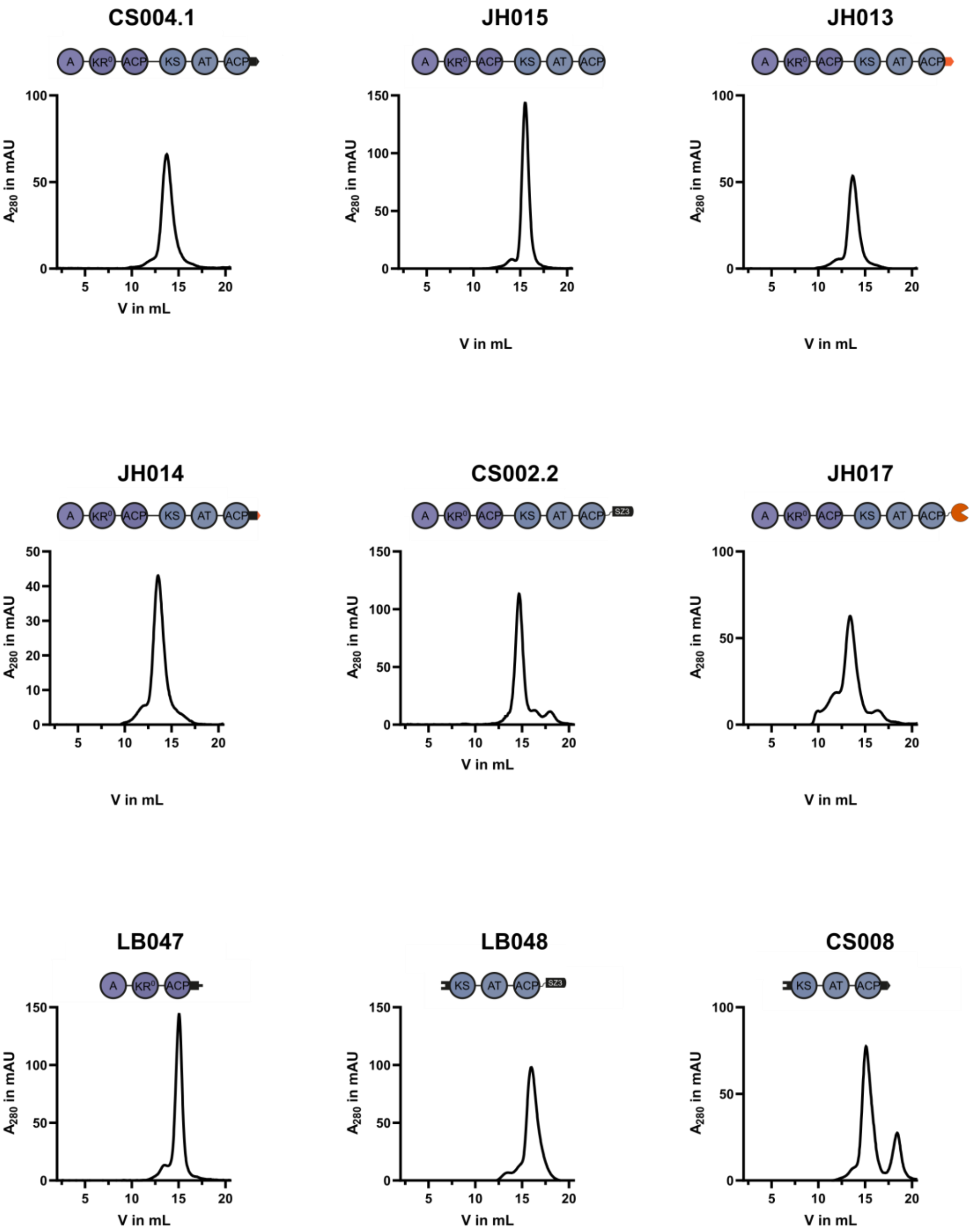

**Figure S2. Analysis of proteins by SEC – VemG and VemG-based constructs.** Size exclusion chromatograms of VemG and VemG-based constructs. All proteins eluted in a predominately single peak from SEC.

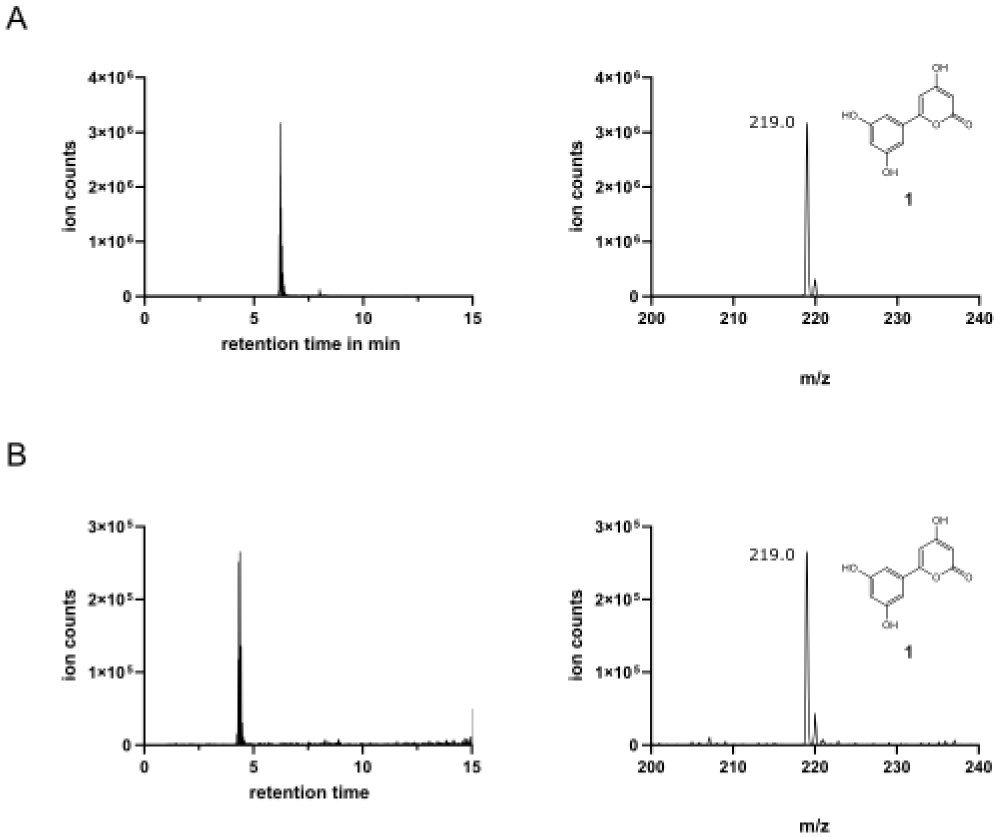

**Figure S3. LC-MS analysis of the venemycin production.** Venemycin ($C_{11}H_8O_5$, exact mass = 220.04 g/mol) could be detected in all reaction mixtures of the HPLC assays after overnight incubation at room temperature. LC-MS analysis is shown for the native VEMS (A) and the DD-less VEMS (B) as an example. The extracted ion chromatograms were obtained by extraction of the $[M\text{-}H]^-$ species.

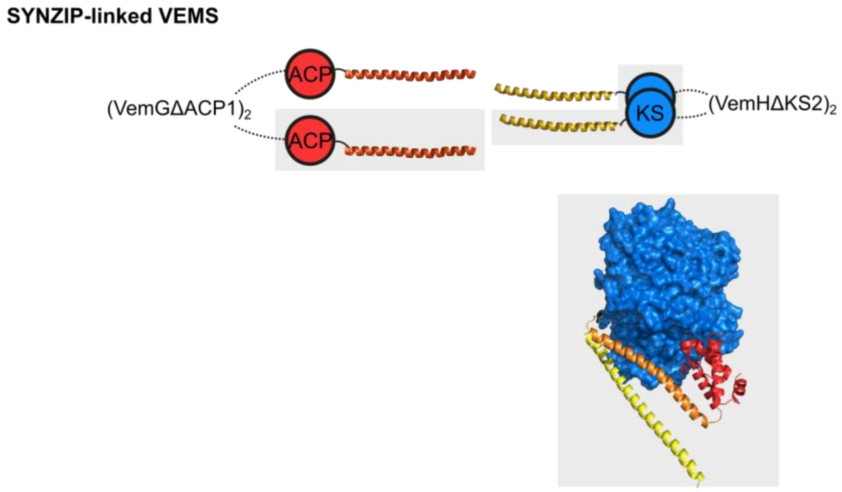

**Figure S4. VemG-SZ3:SZ4-VemH interface predicted with ColabFold[1].** The SYNZIPs were connected to the respective domains by eight-residue $(GGSG)_2$-linkers which are too short to allow the ACP to shuttle to the distant KS active site, if the modules, separated by the rigid SYNZIP interface are linearly orientated. Depicted is the SYNZIP-linked VEMS. The KS2-dimer of VemH is shown as blue surface, ACP1 of VemG is shown in red cartoon, and the interacting SZ3 and SZ4 in orange and yellow, respectively. Parts of the SYNZIP-linked VEMS which are shown in the structure are highlighted in grey. The ACP was predicted to dock to the KS active site entrance at a distance of 14 Å between the cysteine of the KS2 active site and the serine of the ACP1 (phosphopantetheine attachment point), which seems reasonable considering the length of the ~ 18 Å 4'-phosphopantetheine arm[2].

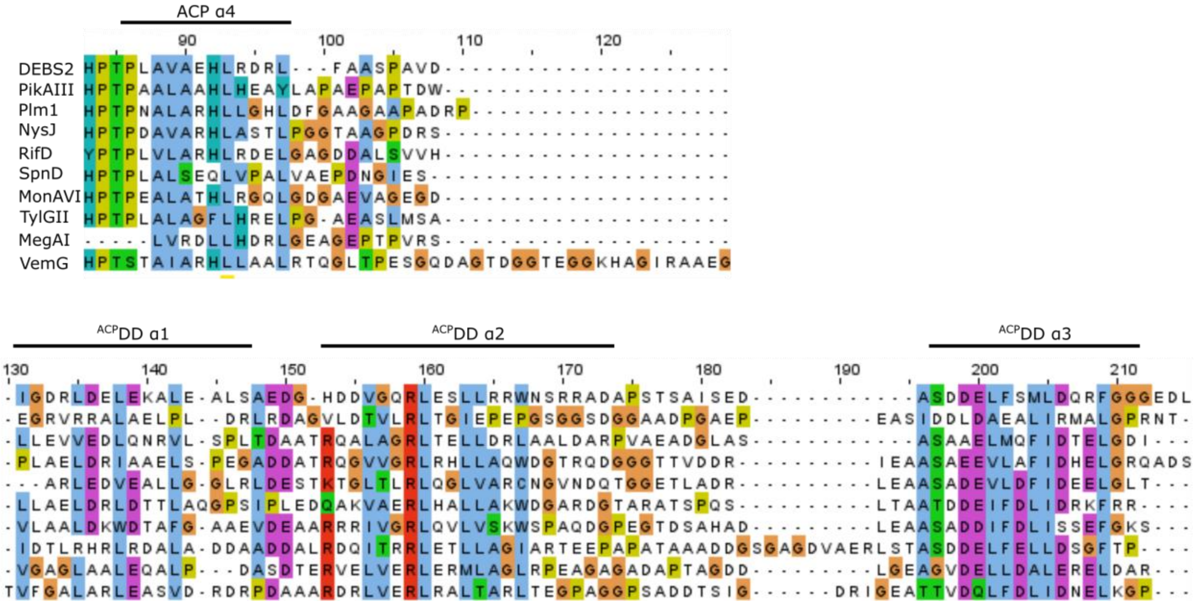

**Figure S5. Alignment of ACP-ACPDD sequences from class 1 *cis*-AT PKS according to Smith *et al.*[3].** Multiple sequence alignment was created with Jalview[4] using Clustal with default settings. Secondary structure elements are indicated as follows: DEBS2 ACP4 α4 according to the DEBS ACP alignment from Alekseyev et al.[5] and the dimerization element (ACPDD α1 + ACPDD α2) and the affinity helix (ACPDD α3) according to the solution structures 1PZQ and 1PZR, respectively. The linker region connecting the ACP and the ACPDD is typically between 8-13 residues long. With a length of 33 residues, the linker connecting the ACP1 and ACPDD of VemG appears to be an exception.

A

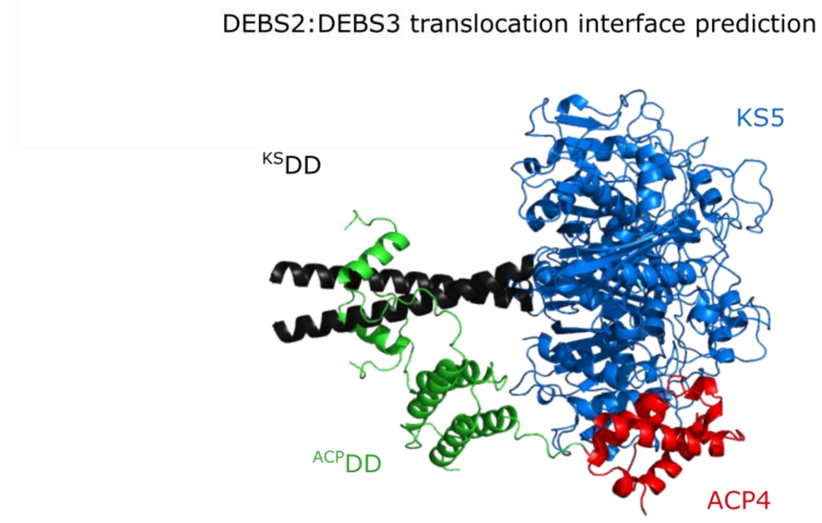

B

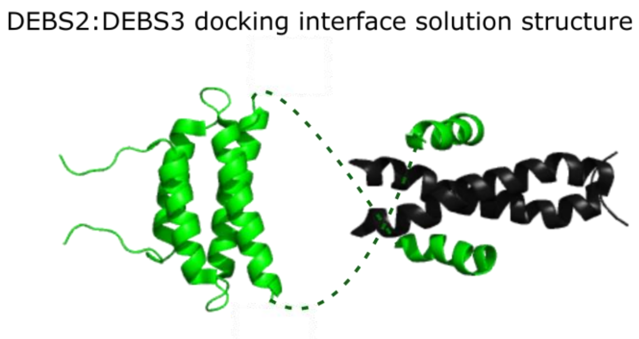

**Figure S6. DEBS2:DEBS3 translocation interface.** (A) DEBS2(ACP4-$^{ACP}$DD) and DEBS3($^{KS}$DD-KS5) complex predicted with ColabFold[1]. The ACP domain was predicted to dock to the KS domain. The distance between the ACP serine (attachment site of the intermediate-bearing phosphopantetheine arm) and the KS active site cysteine is 19.9 Å. The ACP domain can reach the KS entry under reorientation of the docking interface. The flexibility of the docking interface is provided by the linker before and after the dimerization element. (B) DEBS2:DEBS3 docking interface in a parallel orientation according to Broadhurst et al.[6] (PDB ID: 1PZQ and 1PZR). The DEBS2 $^{ACP}$DD is shown in green and the DEBS3 $^{KS}$DD in black. The connection between the dimerization element and the affinity helix of the $^{ACP}$DD is indicated by a dashed line.

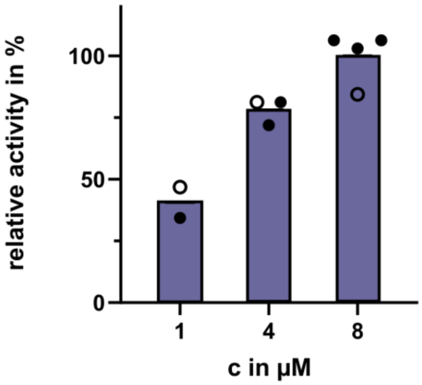

**Figure S7. Activity of the native VEMS at different enzyme concentrations.** Operating the assembly line below the $K_D$ results in a dramatic loss of activity due to dissociation of the VemG:VemH subunit complexes. Assays were performed at 25 °C. Dots of different color represent biological replicates.

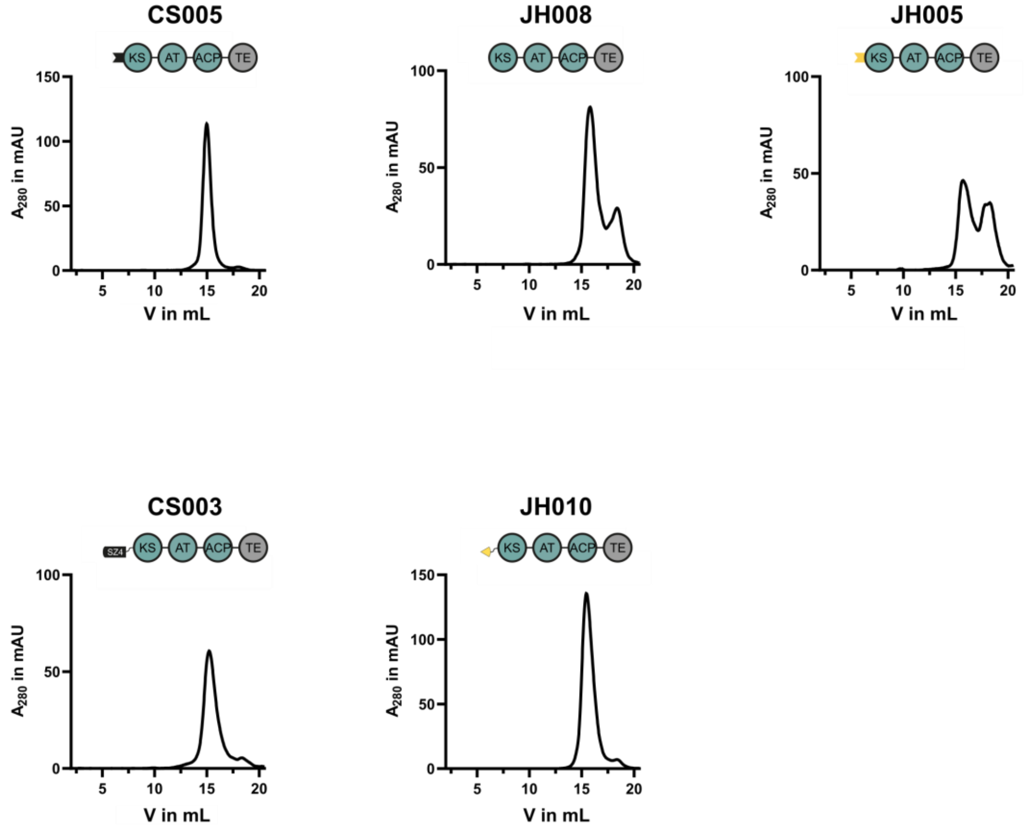

**Figure S8. Analysis of proteins by SEC – VemH and VemH-based constructs.** Size exclusion chromatograms of VemH and VemH-based constructs. All proteins eluted in a predominately single peak from SEC except for JH005 (DEBS (α4)-swapped VemH).

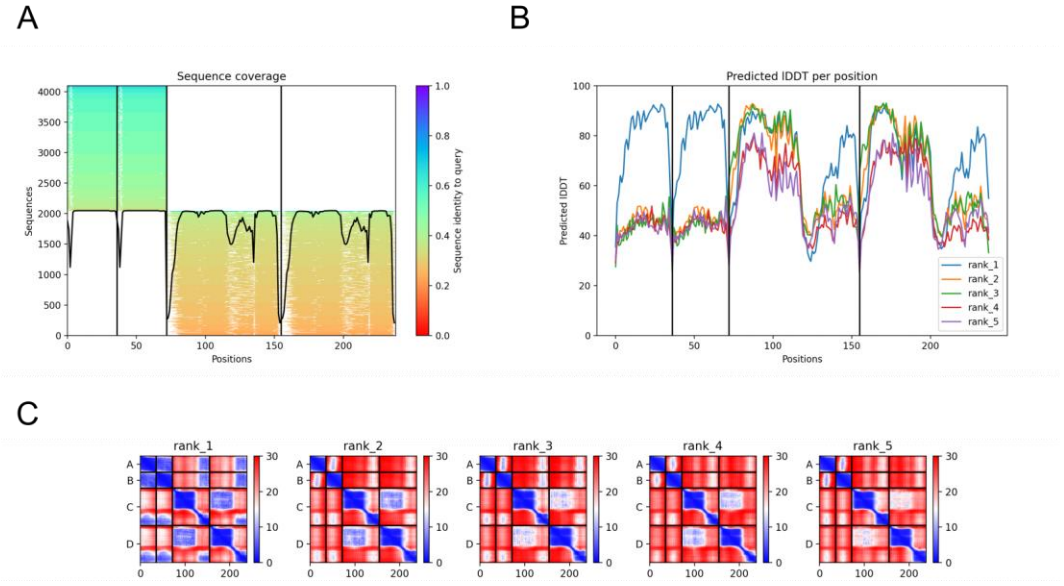

**Figure S9. Quality of the native VemG:VemH docking domain interface predicted with ColabFold[1].** AlphaFold error estimates of the VemG:VemH docking interface shown in Figure 1. (A) Sequence Coverage. (B)Per-residue confidence estimate of predictions (pLDDT) on a scale from 0 – 100 (higher pLDDT is better). (C) Predicted aligned error (PAE) for the five predicted complexes.

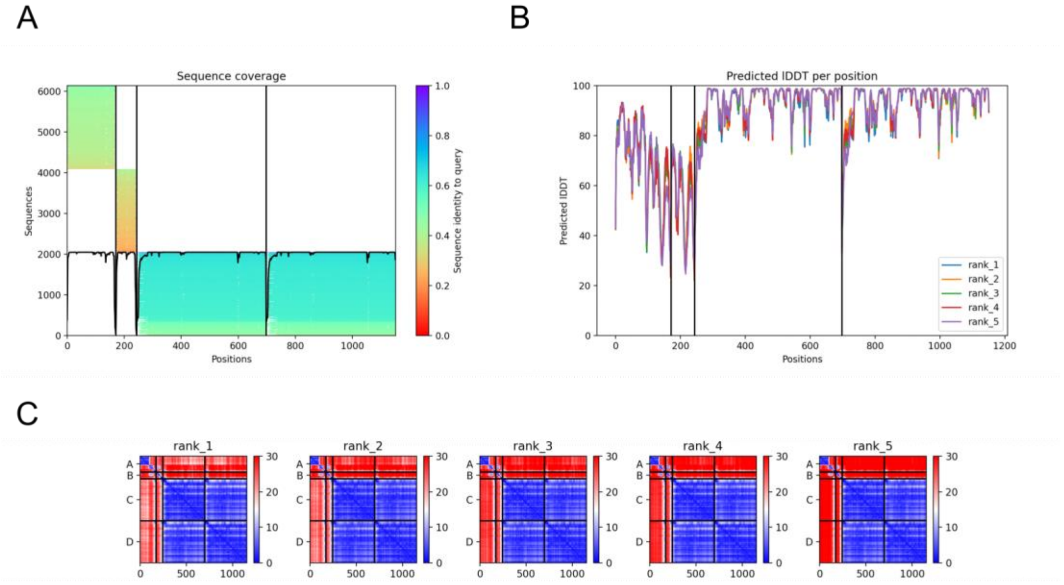

**Figure S10. Quality of the DEBS2:DEBS3 docking domain interface predicted with ColabFold[1].** AlphaFold error estimates of the DEBS2:DEBS3 docking interface shown in Figure S6. (A) Sequence Coverage. (B)Per-residue confidence estimate of predictions (pLDDT) on a scale from 0 – 100 (higher pLDDT is better). (C) Predicted aligned error (PAE) for the five predicted complexes.

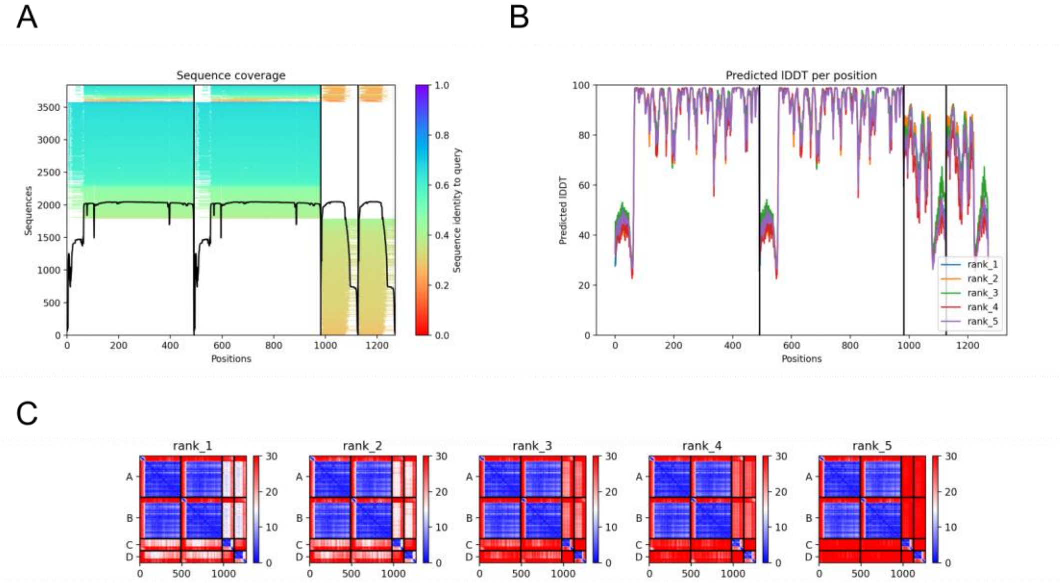

**Figure S11. Quality of the VemG-SZ3:SZ4-VemH interface predicted with ColabFold**[1]. AlphaFold error estimates of the VemG-SZ3:SZ4-VemH docking interface shown in Figure S4. (A) Sequence Coverage. (B)Per-residue confidence estimate of predictions (pLDDT) on a scale from 0 – 100 (higher pLDDT is better). (C) Predicted aligned error (PAE) for the five predicted complexes.

### Supplementary Tables

**Table S1.** Cloning strategy of plasmids generated in this study. Individual fragments were generated by PCR and assembled via In-Fusion cloning. See Table S4 for template sequences.

| Plasmid | Cloning Method | Fragments | Primer Name | Primer Sequence 5'-3' | Template |
|---|---|---|---|---|---|
| pLB047 | In-Fusion | pLB047I | P-LB160 | AAAGGCGCCGGATCCGTGACGCAAGTCGATGACATC | genomic DNA ATCC 10712 |
| | | | P-LB161 | CGGTGAGGCCGCGAAGAGTTCGCCGTCCAGGAC | |
| | | pLB047V | P-LB139 | GGATCCGGCGCCTTTTTCG | pMZ010 |
| | | | P-MZ035 | TTCGCGGCCTCACCGGC | |
| pLB048 | In-Fusion | pLB048I | P-LB162 | GAGCACCGGGCCGGTGACCCGGTCGTCGTCATC | genomic DNA ATCC 10712 |
| | | | P-LB163 | GCCACCGGATCCGCCGAGGGCGGCGAGCAGG | |
| | | pLB048V | P-LB164 | ACCGGCCCGGTGCTCG | pMK150[7] |
| | | | P-MK483 | GGCGGATCCGGTGGCGGA | |
| pCS002.2 | In-Fusion | pCS002.2I | P-CS036 | AGCGCTTGGAGCCATCCAC | pCS004.1 |
| | | | P-CS034 | GATCGCGTCCCGTCCTTCG | |
| | | pCS002.2V | P-CS035 | CGAAGGACGGGACGCGATC | pLB048 |
| | | | P-CS037 | TGTGGATGGCTCCAAGCGCTCATATGTATATCTCCTTCTTAAAGTTAAACAAAATTATTTC | |
| pCS003 | In-Fusion | pCS003I | P-LB166 | ATCCGGTAAGCTTGAGCCGATCGCCATCGTCGG | genomic DNA ATCC 10712 |
| | | | P-LB167 | TGCGGCCGCAAGCTTCAGCAGTGACCGTGACCAC | |
| | | pCS003V | FS15 | CATATGTATATCTCCTTCTTAAAGTTAAACAAAATTATTTC | pMK149[7] |
| | | | P-LB169 | AAGCTTGCGGCCGCACTC | |
| pCS004.1 | In-Fusion | pCS004.1I | P-CS022 | GTGACGCAAGTCGATGACATCC | genomic DNA ATCC |

| Plasmid | Cloning Method | Fragments | Primer Name | Primer Sequence 5'-3' | Template |
|---|---|---|---|---|---|
| | | | P-CS023 | GGGTCCCTTCAGTTCGTTGTC | 10712 |
| | | pCS004.1V | P-CS007 | ATGTCATCGACTTGCGTCACGGATCCGGCGCCTTTTTCG | pLB049 |
| | | | P-CS030 | GAACTGAAGGGACCCCTCGAGCATCATCACCACCAC | |
| pCS005 | Direct transformation of linearized DNA fragment | pCS005 linearized | P-CS024 | TCAAGAAGGTCACCGCCGAACTGCAGGAGACCCGCAGGCAGTTGCGGGGCGCGCTGGCGGCGAGCCGGGAGCCGATCGCCATCGTCG | pLB049 |
| | | | P-CS025 | CGGTGACCTTCTTGAGGTAGTCGACCAGCTTCTCCTCGGTGCCCGTCATATGTATATCTCCTTCTTAAAGTTAAACAAAATTATTTC | |
| pCS008 | In-Fusion | pCS008I | P-CS027 | AAGTCGAGCACCGGGCCGGTGACCCGGTCGTCGTCATCG | genomic DNA ATCC 10712 |
| | | | P-CS026 | TGGTGGTGGTGGTGCTCGAGGGGTCCCTTCAGTTCGTTGTCG | |
| | | pCS008V | P-MZ008 | CTCGAGCACCACCACCAC | pLB048 |
| | | | P-LB164 | ACCGGCCCGGTGCTCG | |
| pJH005 | In-Fusion | pJH005I | P-JH011 | GGGCGACGCTCGACCTGCGTGCCGCCCGGCAGCGCATCCGCGAGCTGGAATCCGAGCCGATCGCCATCGTCG | pCS005 |
| | | | P-LB167 | TGCGGCCGCAAGCTTCAGCAGTGACCGTGACCAC | |
| | | pJH005V | P-LB169 | AAGCTTGCGGCCGCACTC | pCS005 |
| | | | P-JH012 | GGTCGAGCGTCGCCCGACGGAGGTACTCCGCCACCTTCTCGCTGTCAGTCATATGTATATCTCCTTCTTAAAGTTAAACAAAATTATTTC | |
| pJH008 | In-Fusion | pJH008I | P-JH024 | GGAGATATACATATGGAGCCGATCGCCATCGTC | pCS005 |
| | | | P-LB167 | TGCGGCCGCAAGCTTCAGCAGTGACCGTGACCAC | |
| | | pJH008V | P-LB169 | AAGCTTGCGGCCGCACTC | pCS005 |

| Plasmid | Cloning Method | Fragments | Primer Name | Primer Sequence 5'-3' | Template |
|---|---|---|---|---|---|
| | | | P-JH023 | CATATGTATATCTCCTTCTTAAAGTTAAACAAAAT | |
| pJH010 | In-Fusion | pJH010I | P-JH031 | GCAAGCCCTGCGAGCGAGCCGATCGCCATCGTC | pCS005 |
| | | | P-JH032 | AGCAGCCGGATCTCAGTGG | |
| | | pJH010V | P-FB069 | TGAGATCCGGCTGCTAACAAAG | pFB061[8] "spyTseACP" |
| | | | P-MJL014 | GCTCGCAGGGCTTGCCG | |
| pJH013 | In-Fusion | pJH013I | P-LB135 | CTCGAGCATCATCACCACCAC | pCS004.1 |
| | | | P-AR428 | GGATCCGGCGCCTTTTTCG | |
| | | pJH013V | P-LB160 | AAAGGCGCCGGATCCGTGACGCAAGTCGATGACATC | pJH003 |
| | | | P-JH033 | GTGATGATGCTCGAGATCGCCGTCGAGCTCCC | |
| pJH014 | In-Fusion | pJH013I | P-LB135 | CTCGAGCATCATCACCACCAC | pCS004.1 |
| | | | P-AR428 | GGATCCGGCGCCTTTTTCG | |
| | | pJH014V | P-LB160 | AAAGGCGCCGGATCCGTGACGCAAGTCGATGACATC | pJH003 |
| | | | P-JH033 | GTGATGATGCTCGAGATCGCCGTCGAGCTCCC | |
| pJH015 | In-Fusion | pJH013I | P-LB135 | CTCGAGCATCATCACCACCAC | pCS004.1 |
| | | | P-AR428 | GGATCCGGCGCCTTTTTCG | |
| | | pJH015V | P-LB160 | AAAGGCGCCGGATCCGTGACGCAAGTCGATGACATC | pCS004,1 |
| | | | P-JH034 | GTGATGATGCTCGAGCCGGAGGGCGGCGAG | |

| Plasmid | Cloning Method | Fragments | Primer Name | Primer Sequence 5'-3' | Template |
|---|---|---|---|---|---|
| pJH017 | In-Fusion | pJH017I | P-LB160 | AAAGGCGCCGGATCCGTGACGCAAGTCGATGACATC | pCS004.1 |
| | | | P-AR27 | GGTTAGCTCCTTCGGTCCTC | |
| | | pJH017V | P-AR26 | GAGGACCGAAGGAGCTAACC | pJH014 |
| | | | P-AR428 | GGATCCGGCGCCTTTTTCG | |

**Table S2.** DNA and amino acid sequences of plasmids which are not published elsewhere and were used as a template to generate the plasmids of the constructs used in this study. All plasmids were based on a pET22b vector backbone.

| Construct | Sequence form start to stop codon |
|---|---|
| pMZ010 DNA sequence | ATGAGCGCTTGGAGCCATCCACAATTTGAGAAGGGTGGAGGTTCTGGCGGTGGATCGGGAGGTTCAGCGTGGAGCCACCCGCAGTTCGAAAAAGGCGCCGGATCCTCTTCAGCCGGAATTAC<br>CAGGACCGGTGCGAGAACACCGGTGACAGGGCGTGGGGCGGCAGCGTGGGACACGGGGGAAGTGCGGGTCCGACGGGGGTTGCCCCCTGCCGGCCCCGATCATGCGGAGCACTCCTTCTCTC<br>GTGCTCCTACCGGTGATGTGCGCGCCGAATTGATTCGTGGAGAGATGTCGACAGTGTCCAAGAGTGAGTCCGAGGAATTCGTGTCCGTGTCGAACGACGCCGGTTCCGCGCACGGCACAGCG<br>GAACCCGTCGCCGTCGTCGGCATCTCCTGCCGGGTGCCCGGCGCCCGGGACCCGAGAGAGTTCTGGGAACTCCTGGCGGCAGGCGGCCAGGCCGTCACCGACGTCCCCGCGGACCGCTGGAAC<br>GCCGGCGACTTCTACGACCCGGACCGCTCCGCCCCCGGCCGCTCGAACAGCCGGTGGGGCGGGTTCATCGAGGACGTCGACCGGTTCGACGCCGCCTTCTTCGGCATCTCGCCCCGCGAGGCC<br>GCGGAGATGGACCCGCAGCAGCGGCTCGCCCTGGAGCTGGGCTGGGAGGCCCTGGAGCGCGCCGGGATCGACCCGTCCTCGCTCACCGGCACCCGCACCGGCGTCTTCGCCGGCGCCATCTGG<br>GACGACTACGCCACCCTGAAGCACCGCCAGGGCGGCGCCGCGATCACCCCGCACACCGTCACCGGCCTCCACCGCGGCATCATCGCGAACCGACTCTCGTACACGCTCGGGCTCCGCGGCCCCA<br>GCATGGTCGTCGACTCCGGCCAGTCCTCGTCGCTCGTCGCCGTCCACCTCGCGTGCGAGAGCCTGCGGCGCGGCGAGTCCGAGCTCGCCCTCGCCGGCGGCGTCTCGCTCAACCTGGTGCCGG<br>ACAGCATCATCGGGGCGAGCAAGTTCGGCGGCCTCTCCCCCGACGGCCGCGCCTACACCTTCGACGCGCGCGCCAACGGCTACGTACGCGGCGAGGGCGGCGGTTTCGTCGTCCTGAAGCGCC<br>TCTCCCGGGCCGTCGCCGACGGCGACCCGGTGCTCGCCGTGATCCGGGGCAGCGCCGTCAACAACGGCGGCGCCGCCCAGGGCATGACGACCCCCGACGCGCAGGCGCAGGAGGCCGTGCTCC<br>GCGAGGCCCACGAGCGGGCCGGGACCGCGCCGGCCGACGTGCGGTACGTCGAGCTGCACGGCACCGGCACCCCCGTGGGCGACCCGATCGAGGCCGCTGCGCTCGGCGCCGCCCTCGGCACCG<br>GCCGCCCGGCCGGACAGCCGCTCCTGGTCGGCTCGGTCAAGACGAACATCGGCCACCTGGAGGGCGCGGCCGGCATCGCCGGCCTCATCAAGGCCGTCCTGGCGGTCCGCGGTCGCGCGCTGC<br>CCGCCAGCCTGAACTACGAGACCCCGAACCCGGCGATCCCGTTCGAGGAACTGAACCTCCGGGTGAACACGGAGTACCTGCCGTGGGAGCCGGAGCACGACGGGCAGCGGATGGTCGTCGGC<br>GTGTCCTCGTTCGGCATGGGCGGCACGAACGCGCATGTCGTGCTCGAAGAGGCCCCGGGGGTTGTCGAGGGTGCTTCGGTCGTGGAGTCGACGGTCGGCGGGTCGGCGGTCGGCGGCGGTGT<br>GGTGCCGTGGGTGGTGTCGGCGAAGTCCGCTGCCGCGCTGGACGCGCAGATCGAGCGGCTTGCCGCGTTCGCCTCGCGGGATCGTACGGATGGTGTCGACGCGGGCGCTGTCGATGCGGGTG<br>CTGTCGATGCGGGTGCTGTCGCTCGCGTACTGGCCGGCGGGCGTGCTCAGTTCGAGCACCGGGCCGTCGTCGTCGGCAGCGGGCCGGACGATCTGGCGGCAGCGCTGGCCGCGCCTGAGGGTC<br>TGGTCCGGGGCGTGGCTTCCGGTGTCGGGCGAGTGGCGTTCGTGTTCCCCGGGCAGGGCACGCAGTGGGCCGGCATGGGTGCCGAACTGCTGGACTCTTCCGCGGTGTTCGCGGCGGCCATG<br>GCCGAATGCGAGGCCGCACTCTCCCCGTACGTCGACTGGTCGCTGGAGGCCGTCGTACGGCAGGCCCCCGGTGCGCCCACGCTGGAGCGGGTCGATGTCGTGCAGCCTGTGACGTTCGCCGTC<br>ATGGTCTCGCTGGCTCGCGTGTGGCAGCACCACGGGGTGACGCCCCAGGCGGTCGTCGGCCACTCGCAGGGCGAGATCGCCGCCGCGTACGTCGCCGGTGCCCTGAGCCTGGACGACGCCGCT<br>CGTGTCGTGACCCTGCGCAGCAAGTCCATCGCCGCCCACCTCGCCGGCAAGGGCGGCATGCTGTCCCTCGCGCTGAGCGAGGACGCCGTCCTGGAGCGACTGGCCGGGTTCGACGGGCTGTCC<br>GTCGCCGCTGTGAACGGGCCCACCGCCACCGTGGTCTCCGGTGACCCCGTACAGATCGAAGAGCTTGCTCGGGCGTGTGAGGCCGATGGGGTCCGTGCGCGGGTCATTCCCGTCGACTACGCG<br>TCCCACAGCCGGCAGGTCGAGATCATCGAGAGCGAGCTCGCCGAGGTCCTCGCCGGGCTCAGCCCGCAGGCTCCGCGCGTGCCGTTCTTCTCGACACTCGAAGGCGCCTGGATCACCGAGCCC<br>GTGCTCGACGGCGGCTACTGGTACCGCAACCTGCGCCATCGTGTGGGCTTCGCCCCGGCCGTCGAGACCCTGGCCACCGACGAGGGCTTCACCCACTTCGTCGAGGTCAGCGCCCACCCCGTC<br>CTCACCATGGCCCTCCCCGGGACCGTCACCGGTCTGGCGACCCTGCGTCGCGACAACGGCGGTCAGGACCGCCTCGTCGCCTCCCTCGCCGAAGCATGGGCCAACGGACTCGCGGTCGACTGG<br>AGCCCGCTCCTCCCCTCCGCGACCGGCCACCACTCCGACCTCCCCACCTACGCGTTCCAGACCGAGCGCCACTGGCTGGGCGAGATCGAGGCGCTCGCCCCGGCGGGCGAGCCGGCGGTGCAGC<br>CCGCCGTCCTCCGCACGGAGGCGGCCGAGCCGGCGGAGCTCGACCGGGACGAGCAGCTGCGCGTGATCCTGGACAAGGTCCGGGCGCAGACGGCCCAGGTGCTGGGGTACGCGACAGGCGGG<br>CAGATCGAGGTCGACCGGACCTTCCGTGAGGCCGGTTGCACCTCCCTGACCGGCGTGGACCTGCGCAACCGGATCAACGCCGCCTTCGGCGTACGGATGGCGCCGTCCATGATCTTCGACTTC<br>CCCACCCCCGAGGCTCTCGCGGAGCAGCTGCTCCTCGTCGTGCACGGGTTCGCGGCCTCACCGGCGGTGGACATCGGCGACCGGCTGGACGAGCTGGAGAAGGCGCTCGAAGCCCTGTCCGCC<br>GAGGACGGGCACGACGACGTGGGCCAGCGCCTGGAGTCGCTGCTGCGCCGGTGGAACAGCAGGCGGGCGGACGCCCCGAGCACGTCCGCGATCAGCGAGGACGCCAGTGACGACGAGCTGTT<br>CTCGATGCTCGACCAGCGGTTCGGCGGGGGAGAGGACCTGCCGAATTCGAGCTCCGTCGACAAGCTTGCGGCCGCACTCGAGCACCACCACCACCACCACTGA |

| Construct | Sequence form start to stop codon |
|---|---|
| pMZ010 amino acid sequence | MSAWSHPQFEKGGGSGGGSGGSAWSHPQFEKGAGSSSAGITRTGARTPVTGRGAAAWDTGEVRVRRGLPPAGPDHAEHSFSRAPTGDVRAELIRGEMSTVSKSESEEFVSVSNDAGSAHGTA<br>EPVAVVGISCRVPGARDPREFWELLAAGGQAVTDVPADRWNAGDFYDPDRSAPGRSNSRWGGFIEDVDRFDAAFFGISPREAAEMDPQQRLALELGWEALERAGIDPSSLTGTRTGVFAGAI<br>WDDYATLKHRQGGAAITPHTVTGLHRGIIANRLSYTLGLRGPSMVVDSGQSSSLVAVHLACESLRRGESELALAGGVSLNLVPDSIIGASKFGGLSPDGRAYTFDARANGYVRGEGGGFVVLKR<br>LSRAVADGDPVLAVIRGSAVNNGGAAQGMTTPDAQAQEAVLREAHERAGTAPADVRYVELHGTGTPVGDPIEAAALGAALGTGRPAGQPLLVGSVKTNIGHLEGAAGIAGLIKAVLAVRGRA<br>LPASLNYETPNPAIPFEELNLRVNTEYLPWEPEHDGQRMVVGVSSFGMGGTNAHVVLEEAPGVVEGASVVESTVGGSAVGGGVVPWVVSAKSAAALDAQIERLAAFASRDRTDGVDAGAVD<br>AGAVDAGAVARVLAGGRAQFEHRAVVVGSGPDDLAAALAAPEGLVRGVASGVGRVAFVFPGQGTQWAGMGAELLDSSAVFAAAMAECEAALSPYVDWSLEAVVRQAPGAPTLERVDVVQP<br>VTFAVMVSLARVWQHHGVTPQAVVGHSQGEIAAAYVAGALSLDDAARVVTLRSKSIAAHLAGKGGMLSLALSEDAVLERLAGFDGLSVAAVNGPTATVVSGDPVQIEELARACEADGVRARV<br>IPVDYASHSRQVEIIESELAEVLAGLSPQAPRVPFFSTLEGAWITEPVLDGGYWYRNLRHRVGFAPAVETLATDEGFTHFVEVSAHPVLTMALPGTVTGLATLRRDNGGQDRLVASLAEAWA<br>NGLAVDWSPLLPSATGHHSDLPTYAFQTERHWLGEIEALAPAGEPAVQPAVLRTEAAEPAELDRDEQLRVILDKVRAQTAQVLGYATGGQIEVDRTFREAGCTSLTGVDLRNRINAAFGVRM<br>APSMIFDFPTPEALAEQLLLVVHGFAASPAVDIGDRLDELEKALEALSAEDGHDDVGQRLESLLRRWNSRRADAPSTSAISEDASDDELFSMLDQRFGGGEDLPNSSSVDKLAAALEHHHHH<br>H* |
| pLB049 DNA sequence | ATGAGCGCTTGGAGCCATCCACAATTTGAGAAGGGTGGAGGTTCTGGCGGTGGATCGGGAGGTTCAGCGTGGAGCCACCCGCAGTTCGAAAAAGGCGCCGGATCCCAGAAAGTGGCTGAAT<br>TGAAAAACAGAGTTGCTGTTAAACTTAACAGAAATGAACAATTGAAAAACAAGGTAGAAGAGTTGAAAAACCGTAATGCTTACCTGAAAAACGAACTGGCTACATTAGAAAATGAAGTCG<br>CCAGATTGGAGAACGATGTTGCTGAAGGCGGATCCGGTGGCGGATCCGGTAAGCTTGAGCCGATCGCCATCGTCGGCATGGCCTGCCGCTACCCCGGCGGCGTCCGGACGCCCGAGGCCCTG<br>TGGCGGCTCGTCCTGGACGAACAGGACGCCATCTCCGGCTTCCCGACGAACCGGGGCTGGGACATCGACGGCATCTACCACCCCGACCCCGACCGGCCCGGCACGTGCTACGCCCGGGAGGGC<br>GGCTTCCTCCACGACGCCGCCCTGTTCGACGCGGAGTTCTTCGGGGTGTCGCCCCGCGAGGCCCAGGCGATGGACCCGCAGCAGCGGCTCCTCCTGGAGACCGCCTGGGAGGCGTTCGAGCGG<br>GCCGGGATCGACCCGACCTCGCTGCGCGGCAGCGACACGGGCGTGTTCGCCGGGGTCGTCCACCACGACTACGCGACGGCCCGGGTGCCGGAGACACTGGAGCCCTATCTCGTCACGGGCCTC<br>TCCGGCGGCGTGGCGTCGGGCCGCATCGCGTACACCTTCGGCTTCGAGGGTCCCGCGGTCACCGTGGACACCGCCTGCTCCTCGTCCCTCGTCGCCCTGCACCAGGCGGCGCACGCACTGCGCT<br>CCGGAGAGTGCGAGCTGGCCCTCGCGGGCGGTGTGACGATCATGGCGACCCCGCGCGCCTTCCTCTCCTTCAGCCGGCAGCGCGGCCTGTCGCCCGACGGACGCTGCCGGGCGTTCGGCGCGG<br>GCGCCGACGGCACCGGCTGGGCCGAGGGCGCGGGCATGCTGCTCGTCGAGCGGCTGTCCGACGCCCGGCGCAAGGGCCACCCGGTGCTCGCCGTCCTCCGCGGCTCCGCGGTCAACCAGGACG<br>GCGCCTCCAACGGCCTCTCCGCCCCCAACGGCCCCTCGCAGCAACGGGTCATCCGCAAGGCCCTGGCCCACGCGGGCCTCGCCGCCCGGGACGTCGACGTCGTCGAG GGCACGGGACCGGGA<br>CGAAGCTCGGCGACCCGATCGAGGCGCAGGCGCTCCTCGCCACCTACGGCCAGGAGCGCGACGCGGGCCTCCCGCTCCACCTCGGCTCGATGAAGTCGAACGTGGGCCACTCCCAGGCGGCGG<br>CCGGGGTCGGCGGCGTGATCAAGATGATCGAGGCGATGCGGCACGGCATCCTGCCGAGGACCCTGCACGCCGACGAGCCGACCCCGCACGTCGACTGGTCGGCCGGCGACATCGAGCTCCTGA<br>CCCGGCGGCGCGCCTGGCCCGAGACGGGACGCCCGCGCCGGGCGGCGGTGTCGTCGTTCGGCATCAGCGGCACCAACGCCCACGTGATCCTGGAGGCCCCACCCGAGGAGCCCGCCAGGGACG<br>CCGCGGAGACCCGCCCCGAGGCGCGGTCGCGGGAGGCCGCGGAGGGCCGCCGCGAGGCCGCCGGGCAGGGCCGGACGGAGACCGGGCCCGGGCCGTCCGCGACCCCTCCCGAGGCGCCCGGCC<br>GCGCCCGGCCGCCGGTGCCCTGGCCGCTCTCCGGGCGGGACGCCGGAGCGCTGCGGGACCAGATCGGGCGGCTGCGGGCCCATCTCGACGCGGCTCCCGCCGACCCCGAGGACGTGGCCCACT<br>CGCTCGCGCGGCGCGCGGTCTTCCGGCACCGGGCGGTGCTCCTCGCGGCGCCGCAGGCGCCGGCGGGCGGCTCCCCCCGCGCGGTCACCGGCGTCGCGCGCCCCGGCGGCACCGCGCTGCTCTT<br>CTCCGGCCAGGGCAGCCAACGCGTCGGCATGGGCAGTGAGTTGTACGAGACGTACCCGGTCTTCGCGGAGTCCTTCGACGCCGTCGCGGAGCACACCGGACTGCCCCTGAAGGACGTGGTCCT<br>CGGCGGGACCCCGGACGGCCTCCTCGACCGCACGCGCTACGCCCAGCCCGCGCTGTTCGCCGTCGAGGTGTCCCTGTTCCGCCTGGTGCGCGCCCTGGGCCTCGACGTCCGGGCCGTGGTCGGC<br>CACTCGGTGGGCGAGATCGCCGCCGCCCACGTCGCCGGAGTGATGTCCATGGCGGACGCCTGCCGCCTGGTGGAGGCGCGCGGACGGCTGATGGACGCGCTGCCGCCGGGCGGCGCCATGGTG<br>GCCGTGGAGGTCACCGAGGCCGAGGCCTCGGCTGCCCTGGCCGGCCTGGAGGACCGGGTCGCCGTCGCCGCCGTCAACGGTCCCGCGTCCACGGTGCTCTCCGGCGAGGAGGGCGCCGTCCTC<br>AAGCTGGCCGACGCCTGGCGTGAACGCGGCGTCCGCACGCACCGGCTGACGGTCAGTCACGCCTTTCACTCGCCGCTCATGGAGCCCATGGTCGACGCCTTCCGCGAGGTCGTGGCGGGCCTG<br>GACCTCCACCGGCCCACTCTGGCCGGACTGCCCGCCGAGGTCGTCGACCCCGAGTACTGGGTGCGGCACGTCCGCCGCCCGGTGCGGTTCGCCGACGCCGTGGCGCGGGCCCGTGAGGCCGGC<br>GCGGTCCGCTGGCTGGAGGTGGGGCCGGGCGGCGTCCTGACCGCCCTGGCCCAGCGGATCGTGCCCGACACCGAGGAGCACGTCTTCGCCGCGGCGCTGCGGACGGACCGGCCGGAACCCGAG<br>GCGCTCCTCGTGGCCCTGTCCCAGGTCCACGTGGACGGCGGCACGGTCGACTGGAGCGGGCTCTGTGCCGGGGGCAGGCTGGTGGACCTCCCCACGTACCCCTTCCAGCGACAGCACTACTGG<br>ATCGAGGACCAGCCCCTGCCGCCGACCGCGCCCAGACCGGGGACCGCCCCGTCCGGCACCGGGACCGCGGCGGAGGGGGCGGCGGCCGCCGAGGTCCCCCTCTCCGAGCGGCTCGCCCGGCTG<br>ACCGGCGCGGAACGGCTCGCCGCCGTGCGCGAACTCGTCCTCGCCGAGGCCTCGGAGACCCTCGGCCACACCGGCACCCTGATCACCGCCGACCGCACCCGGCAGGAGCTGGGCTTCGACTCG<br>CTCACCGCGATCGAACTCCGCAACCGGATCAGCCGGACCCTCGGCGTCCGCCTCCCCCCGACGCTCGTCTTCGACCACGAGGACCTGGGCGAGATCGCCTCCTTCGTCGACGCCCGCCTCGACG<br>ACGCCGCCACCGGCCGGTCCACCGGCCACGGCCCCCTCGGTGAGGACGGCTCCGGCCTGCTCACGGAGCTCTTCAGGGAGGCGGCGGCCGCCGGCCGGCTCGACGACGCCGTGACCCTCACGG<br>AGGCCGCCGCCCGGATGCGGCGGACCTTCACCGACGCCGAGGACCCGGCGGTCCGGCGCACCCCGGTCTGGTTCGGCCGGGGGCCGGCCCGCCCCACGGTCGTCTGCCTGCCGTCGTTCAGCGC<br>GATCGCCGGCGTCCACGTGTACGCCCGGTTCGCGGACGCCTTCGGCGACGGCTGGCGGGTGGCCGCGCTGGCCCATCCGGGCTTCGTCCCCGGCGAGCCGCTGCCGGACTCCGTCGACGTCCT<br>GGCCGAGCTGCACGCCCGCACGGTGCTCGACACGGTCGGCGCGGACCCGTTCCTCCTGGTCGGACGCTCCGCCGGCGGCTGGGTGGCCCACGAGGTGGCGGCCGTCCTGGAGCGGATGGGCAG<br>GGCGCCGGACGGTGTCGCCCTGCTCGACACCCCGGCCCGCGCCGACGACCCGCGCGGGCACGCGGTGATGGTCGGCGGCATGCTGGAGCGCGACAGCCGGCTCGTCACGATCGACGACTACCG<br>GCTGACCGCGATGGGCGGCTACTCCCGGCTGTTCCGGGAGTGGAAGCCCGAACCGATCGCCGCCGCCACCCTGCTGGTGCACGCCGCCACCCCCTACGGGGCCGACGAGGCGCGGATCGCCTC<br>CTGGGACCTCCCGCACCAGGCGGTCAAGGTGACGGGCGATCACTTCACCATGCTGGAGCGGCACTCGGCGACGACGGCCGAGGCGGTCGAGCAGTGGTCACGGTCACTGCTGAAGCTTGCGG<br>CCGCACTCGAGCATCATCACCACCACCACCACCACTGA |

| Construct | Sequence form start to stop codon |
|---|---|
| pLB049 amino acid sequence | MSAWSHPQFEKGGGSGGGSGGSAWSHPQFEKGAGSQKVAELKNRVAVKLNRNEQLKNKVEELKNRNAYLKNELATLENEVARLENDVAEGGSGGGSGKLEPIAIVGMACRYPGGVRTPEAL<br>WRLVLDEQDAISGFPTNRGWDIDGIYHPDPDRPGTCYAREGGFLHDAALFDAEFFGVSPREAQAMDPQQRLLLETAWEAFERAGIDPTSLRGSDTGVFAGVVHHDYATARVPETLEPYLVT<br>GLSGGVASGRIAYTFGFEGPAVTVDTACSSSLVALHQAAHALRSGECELALAGGVTIMATPRAFLSFSRQRGLSPDGRCRAFGAGADGTGWAEGAGMLLVERLSDARRKGHPVLAVLRGSAVN<br>QDGASNGLSAPNGPSQQRVIRKALAHAGLAARDVDVVEGHGTGTKLGDPIEAQALLATYGQERDAGLPLHLGSMKSNVGHSQAAAGVGGVIKMIEAMRHGILPRTLHADEPTPHVDWSAGD<br>IELLTRRRAWPETGRPRRAAVSSFGISGTNAHVILEAPPEEPARDAAETRPEARSREAAEGRREAAGQGRTETGPGPSATPPEAPGRARPPVPWPLSGRDAGALRDQIGRLRAHLDAAPADPE<br>DVAHSLARRAVFRHRAVLLAAPQAPAGGSPRAVTGVARPGGTALLFSGQGSQRVGMGSELYETYPVFAESFDAVAEHTGLPLKDVVLGGTPDGLLDRTRYAQPALFAVEVSLFRLVRALGLD<br>VRAVVGHSVGEIAAAHVAGVMSMADACRLVEARGRLMDALPPGGAMVAVEVTEAEASAALAGLEDRVAVAAVNGPASTVLSGEEGAVLKLADAWRERGVRTHRLTVSHAFHSPLMEPMV<br>DAFREVVAGLDLHRPTLAGLPAEVVDPEYWVRHVRRPVRFADAVARAREAGAVRWLEVGPGGVLTALAQRIVPDTEEHVFAAALRTDRPEPEALLVALSQVHVDGGTVDWSGLCAGGRLV<br>DLPTYPFQRQHYWIEDQPLPPTAPRPGTAPSGTGTAAEGAAAAEVPLSERLARLTGAERLAAVRELVLAEASETLGHTGTLITADRTRQELGFDSLTAIELRNRISRTLGVRLPPTLVFDHEDL<br>GEIASFVDARLDDAATGRSTGHGPLGEDGSGLLTELFREAAAAGRLDDAVTLTEAAARMRRTFTDAEDPAVRRTPVWFGRGPARPTVVCLPSFSAIAGVHVYARFADAFGDGWRVAALAHP<br>GFVPGEPLPDSVDVLAELHARTVLDTVGADPFLLVGRSAGGWVAHEVAAVLERMGRAPDGVALLDTPARADDPRGHAVMVGGMLERDSRLVTIDDYRLTAMGGYSRLFREWKPEPIAAAT<br>LLVHAATPYGADEARIASWDLPHQAVKVTGDHFTMLERHSATTAEAVEQWSRSLLKLAAALEHHHHHHHH* |
| pJH003 DNA sequence | ATGAGCGCTTGGAGCCATCCACAATTTGAGAAGGGTGGAGGTTCTGGCGGTGGATCGGGAGGTTCAGCGTGGAGCCACCCGCAGTTCGAAAAAGGCGCCGGATCCCAGAAAGTGGCTGAAT<br>TGAAAAACAGAGTTGCTGTTAAACTTAACAGAAATGAACAATTGAAAAACAAGGTAGAAGAGTTGAAAAACCGTAATGCTTACCTGAAAAACGAACTGGCTACATTAGAAAATGAAGTCG<br>CCAGATTGGAGAACGATGTTGCTGAAGGCGGATCCGGTGGCGGATCCGGTAAGCTTGAGCCGATCGCCATCGTCGGCATGGCCTGCCGCTACCCCGGCGGCGTCCGGACGCCCGAGGCCCTG<br>TGGCGGCTCGTCCTGGACGAACAGGACGCCATCTCCGGCTTCCCGACGAACCGGGGCTGGGACATCGACGGCATCTACCACCCCGACCCCGACCGGCCCGGCACGTGCTACGCCCGGGAGGGC<br>GGCTTCCTCCACGACGCCGCCCTGTTCGACGCGGAGTTCTTCGGGGTGTCGCCCCGCGAGGCCCAGGCGATGGACCCGCAGCAGCGGCTCCTCCTGGAGACCGCCTGGGAGGCGTTCGAGCGG<br>GCCGGGATCGACCCGACCTCGCTGCGCGGCAGCGACACGGGCGTGTTCGCCGGGGTCGTCCACCACGACTACGCGACGGCCCGGGTGCCGGAGACACTGGAGCCCTATCTCGTCACGGGCCTC<br>TCCGGCGGCGTGGCGTCGGGCCGCATCGCGTACACCTTCGGCTTCGAGGGTCCCGCGGTCACCGTGGACACCGCCTGCTCCTCGTCCCTCGTCGCCCTGCACCAGGCGGCGCACGCACTGCGCT<br>CCGGAGAGTGCGAGCTGGCCCTCGCGGGCGGTGTGACGATCATGGCGACCCCGCGCGCCTTCCTCTCCTTCAGCCGGCAGCGCGGCCTGTCGCCCGACGGACGCTGCCGGGCGTTCGGCGCGG<br>GCGCCGACGGCACCGGCTGGGCCGAGGGCGCGGGCATGCTGCTCGTCGAGCGGCTGTCCGACGCCCGGCGCAAGGGCCACCCGGTGCTCGCCGTCCTCCGCGGCTCCGCGGTCAACCAGGACG<br>GCGCCTCCAACGGCCTCTCCGCCCCCAACGGCCCCTCGCAGCAACGGGTCATCCGCAAGGCCCTGGCCCACGCGGGCCTCGCCGCCCGGGACGTCGACGTCGTCGAGGGGCACGGGACCGGGA<br>CGAAGCTCGGCGACCCGATCGAGGCGCAGGCGCTCCTCGCCACCTACGGCCAGGAGCGCGACGCGGGCCTCCCGCTCCACCTCGGCTCGATGAAGTCGAACGTGGGCCACTCCCAGGCGGCGG<br>CCGGGGTCGGCGGCGTGATCAAGATGATCGAGGCGATGCGGCACGGCATCCTGCCGAGGACCCTGCACGCCGACGAGCCGACCCCGCACGTCGACTGGTCGGCCGGCGACATCGAGCTCCTGA<br>CCCGGCGGCGCGCCTGGCCCGAGACGGGACGCCCGCGCCGGGCGGCGGTGTCGTCGTTCGGCATCAGCGGCACCAACGCCCACGTGATCCTGGAGGCCCCACCCGAGGAGCCCGCCAGGGACG<br>CCGCGGAGACCCGCCCCGAGGCGCGGTCGCGGGAGGCCGCGGAGGGCCGCCGCGAGGCCGCCGGGCAGGGCCGGACGGAGACCGGGCCCGGGCCGTCCGCGACCCCTCCCGAGGCGCCCGGCC<br>GCGCCCGGCCGCCGGTGCCCTGGCCGCTCTCCGGGCGGGACGCCGGAGCGCTGCGGGACCAGATCGGGCGGCTGCGGGCCCATCTCGACGCGGCTCCCGCCGACCCCGAGGACGTGGCCCACT<br>CGCTCGCGCGGCGCGCGGTCTTCCGGCACCGGGCGGTGCTCCTCGCGGCGCCGCAGGCGCCGGCGGGCGGCTCCCCCCGCGCGGTCACCGGCGTCGCGCGCCCCGGCGGCACCGCGCTGCTCTT<br>CTCCGGCCAGGGCAGCCAACGCGTCGGCATGGGCAGTGAGTTGTACGAGACGTACCCGGTCTTCGCGGAGTCCTTCGACGCCGTCGCGGAGCACACCGGACTGCCCCTGAAGGACGTGGTCCT<br>CGGCGGGACCCCGGACGGCCTCCTCGACCGCACGCGCTACGCCCAGCCCGCGCTGTTCGCCGTCGAGGTGTCCCTGTTCCGCCTGGTGCGCGCCCTGGGCCTCGACGTCCGGGCCGTGGTCGGC<br>CACTCGGTGGGCGAGATCGCCGCCGCCCACGTCGCCGGAGTGATGTCCATGGCGGACGCCTGCCGCCTGGTGGAGGCGCGCGGACGGCTGATGGACGCGCTGCCGCCGGGCGGCGCCATGGTG<br>GCCGTGGAGGTCACCGAGGCCGAGGCCTCGGCTGCCCTGGCCGGCCTGGAGGACCGGGTCGCCGTCGCCGCCGTCAACGGTCCCGCGTCCACGGTGCTCTCCGGCGAGGAGGGCGCCGTCCTC<br>AAGCTGGCCGACGCCTGGCGTGAACGCGGCGTCCGCACGCACCGGCTGACGGTCAGTCACGCCTTTCACTCGCCGCTCATGGAGCCCATGGTCGACGCCTTCCGCGAGGTCGTGGCGGGCCTG<br>GACCTCCACCGGCCCACTCTGGCCGGACTGCCCGCCGAGGTCGTCGACCCCGAGTACTGGGTGCGGCACGTCCGCCGCCCGGTGCGGTTCGCCGACGCCGTGGCGCGGGCCCGTGAGGCCGGC<br>GCGGTCCGCTGGCTGGAGGTGGGGCCGGGCGGCGTCCTGACCGCCCTGGCCCAGCGGATCGTGCCCGACACCGAGGAGCACGTCTTCGCCGCGGCGCTGCGGACGGACCGGCCGGAACCCGAG<br>GCGCTCCTCGTGGCCCTGTCCCAGGTCCACGTGGACGGCGGCACGGTCGACTGGAGCGGGCTCTGTGCCGGGGGCAGGCTGGTGGACCTCCCCACGTACCCCTTCCAGCGACAGCACTACTGG<br>ATCGAGGACCAGCCCCTGCCGCCGACCGCGCCCAGACCGGGGACCGCCCCGTCCGGCACCGGGACCGCGGCGGAGGGGGCGGCGGCCGCCGAGGTCCCCCTCTCCGAGCGGCTCGCCCGGCTG<br>ACCGGCGCGGAACGGCTCGCCGCCGTGCGCGAACTCGTCCTCGCCGAGGCCTCGGAGACCCTCGGCCACACCGGCACCCTGATCACCGCCGACCGCACCCGGCAGGAGCTGGGCTTCGACTCG<br>CTCACCGCGATCGAACTCCGCAACCGGATCAGCCGGACCCTCGGCGTCCGCCTCCCCCCGACGCTCGTCTTCGACCACGAGGACCTGGGCGAGATCGCCTCCTTCGTCGACGCCCGCCTCGACG<br>ACGCCGCCACCGGCCGGTCCACCGGCCACGGCCCCCTCGGTGAGGACGGCTCCGGCCTGCTCACGGAGCTCTTCAGGGAGGCGGCGGCCGCCGGCCGGCTCGACGACGCCGTGACCCTCACGG<br>AGGCCGCCGCCCGGATGCGGCGGACCTTCACCGACGCCGAGGACCCGGCGGTCCGGCGCACCCCGGTCTGGTTCGGCCGGGGGCCGGCCCGCCCCACGGTCGTCTGCCTGCCGTCGTTCAGCGC<br>GATCGCCGGCGTCCACGTGTACGCCCGGTTCGCGGACGCCTTCGGCGACGGCTGGCGGGTGGCCGCGCTGGCCCATCCGGGCTTCGTCCCCGGCGAGCCGCTGCCGGACTCCGTCGACGTCCT<br>GGCCGAGCTGCACGCCCGCACGGTGCTCGACACGGTCGGCGCGGACCCGTTCCTCCTGGTCGGACGCTCCGCCGGCGGCTGGGTGGCCCACGAGGTGGCGGCCGTCCTGGAGCGGATGGGCAG<br>GGCGCCGGACGGTGTCGCCCTGCTCGACACCCCGGCCCGCGCCGACGACCCGCGCGGGCACGCGGTGATGGTCGGCGGCATGCTGGAGCGCGACAGCCGGCTCGTCACGATCGACGACTACCG<br>GCTGACCGCGATGGGCGGCTACTCCCGGCTGTTCCGGGAGTGGAAGCCCGAACCGATCGCCGCCGCCACCCTGCTGGTGCACGCCGCCACCCCCTACGGGGCCGACGAGGCGCGGATCGCCTC<br>CTGGGACCTCCCGCACCAGGCGGTCAAGGTGACGGGCGATCACTTCACCATGCTGGAGCGGCACTCGGCGACGACGGCCGAGGCGGTCGAGCAGTGGTCACGGTCACTGCTGAAGCTTGCGG<br>CCGCACTCGAGCATCATCACCACCACCACCACCACTGA |

| Construct | Sequence form start to stop codon |
|---|---|
| pJH003 amino acid sequence | MSAWSHPQFEKGGGSGGGSGGSAWSHPQFEKGAGSQKVAELKNRVAVKLNRNEQLKNKVEELKNRNAYLKNELATLENEVARLENDVAEGGSGGGSGKLEPIAIVGMACRYPGGVRTPEAL<br>WRLVLDEQDAISGFPTNRGWDIDGIYHPDPDRPGTCYAREGGFLHDAALFDAEFFGVSPREAQAMDPQQRLLLETAWEAFERAGIDPTSLRGSDTGVFAGVVHHDYATARVPETLEPYLVT<br>GLSGGVASGRIAYTFGFEGPAVTVDTACSSSLVALHQAAHALRSGECELALAGGVTIMATPRAFLSFSRQRGLSPDGRCRAFGAGADGTGWAEGAGMLLVERLSDARRKGHPVLAVLRGSAVN<br>QDGASNGLSAPNGPSQQRVIRKALAHAGLAARDVDVVEGHGTGTKLGDPIEAQALLATYGQERDAGLPLHLGSMKSNVGHSQAAAGVGGVIKMIEAMRHGILPRTLHADEPTPHVDWSAGD<br>IELLTRRRAWPETGRPRRAAVSSFGISGTNAHVILEAPPEEPARDAAETRPEARSREAAEGRREAAGQGRTETGPGPSATPPEAPGRARPPVPWPLSGRDAGALRDQIGRLRAHLDAAPADPE<br>DVAHSLARRAVFRHRAVLLAAPQAPAGGSPRAVTGVARPGGTALLFSGQGSQRVGMGSELYETYPVFAESFDAVAEHTGLPLKDVVLGGTPDGLLDRTRYAQPALFAVEVSLFRLVRALGLD<br>VRAVVGHSVGEIAAAHVAGVMSMADACRLVEARGRLMDALPPGGAMVAVEVTEAEASAALAGLEDRVAVAAVNGPASTVLSGEEGAVLKLADAWRERGVRTHRLTVSHAFHSPLMEPMV<br>DAFREVVAGLDLHRPTLAGLPAEVVDPEYWVRHVRRPVRFADAVARAREAGAVRWLEVGPGGVLTALAQRIVPDTEEHVFAAALRTDRPEPEALLVALSQVHVDGGTVDWSGLCAGGRLV<br>DLPTYPFQRQHYWIEDQPLPPTAPRPGTAPSGTGTAAEGAAAAEVPLSERLARLTGAERLAAVRELVLAEASETLGHTGTLITADRTRQELGFDSLTAIELRNRISRTLGVRLPPTLVFDHEDL<br>GEIASFVDARLDDAATGRSTGHGPLGEDGSGLLTELFREAAAAGRLDDAVTLTEAAARMRRTFTDAEDPAVRRTPVWFGRGPARPTVVCLPSFSAIAGVHVYARFADAFGDGWRVAALAHP<br>GFVPGEPLPDSVDVLAELHARTVLDTVGADPFLLVGRSAGGWVAHEVAAVLERMGRAPDGVALLDTPARADDPRGHAVMVGGMLERDSRLVTIDDYRLTAMGGYSRLFREWKPEPIAAAT<br>LLVHAATPYGADEARIASWDLPHQAVKVTGDHFTMLERHSATTAEAVEQWSRSLLKLAAALEHHHHHHHH* |

**Table S3.** Plasmids used in this study and their origin

| Plasmid | Origin |
|---|---|
| pLB047_twinstrep_VemG(M0)-DEBS(DD4)-H6_pET22b_carb | this study |
| pLB048_DEBS(DD5)-VemG(M1)-(GGSG)$_2$-SZ3-H6_pET22b_carb | this study |
| pCS002.2_twinstrep-VemG(M0-M1)-(GGSG)$_2$-SZ3-H6_pET22b_carb | this study |
| pCS003_SZ4-(GGSG)$_2$-VemH(M2-TE)-H8_pET22b_carb | this study |
| pCS004.1_twinstrep-VemG(M0-M1-DD)-H8_pET22b_carb | this study |
| pCS005_VemH(DD2-M2-TE)-H8_pET22b_carb | this study |
| pCS008_DEBS(DD5)-VemG(M1-DD)-H6_pET22b_carb | this study |
| pJH005_DEBS(DD3)-VemH(M2-TE)-H8_pET22b_carb | this study |
| pJH008_VemH(noDD-M2-TE)-H8_pET22b_carb | this study |
| pJH010_SpyT-PAASlinker-VemH(M2-TE)-H8_pET22b_carb | this study |
| pJH013_twinstrep-VemG(M0-M1)-DEBS(DD2a1-a3)-H8_pET22b_carb | this study |
| pJH014_twinstrep-VemG(M0-M1)-DEBS(DD2a3)-H8_pET22b_carb | this study |
| pJH015_twinstrep-VemG(M0-M1-noDD)-H8_pET22b_carb | this study |
| pJH017_twinStrep-VemG(M0-M1-a1-a2)-SpyC-H8_pET22b_carb | this study |
| pAR357_SFP_pCDF-1b | Rittner *et al.*[9] |

**Table S4.** Amino acid sequences of VemG and VemG-based constructs

| Construct | Amino acid sequence |
|---|---|
| VemG (M0-M1-DD1) CS004.1 | MSAWSHPQFEKGGGSGGGSGGSAWSHPQFEKGAGSVTQVDDIRPLSAVLRAHALDRPDKVAFADEERHVTYARLAERTGRLAGHLAG HGLRRGDRVAILLGNSVTTVESYLAVTRAAAVGVPVNPQSSDAELAHQLDDSGARFVITDGPHLDQVERLRATREGIRVVLARGYGP GGGPHGPGPEAPASEGAPALDGTSAPDGAPGPDGPLSFEELAGSEPAQAPRDDLGLDEPAWILYTSGTTGAAKGVVSHQRACLWSVR SSYQGVLGLGPDDRLLWPLPLFHSLAHILCVLGVTVSGATARILPGFGARDVLDALRAEPCTMLLGVPSMYLRLVAAVEAVEEAGGS VPKPRLCVVTGAATGPELASAVERVLGAPLVNSYGATETCGAITLSRPDSERPPGTVGTAVPGSELRLVDPRTGRDARTGDEGEVLV RSPGLLLRYHDRPEETAAALRDGWYRTGDLARRDAAGRLTITGRVKELVIRAGENIQPGEIEDVLRTVPGIADAAVTGRPDEALGEV PVAYVVPAAGGWSPAEALAACRERLSYFKVPAELYEIAALPVTGSGKVTRRALPGLPARLRALGTGHHERLWRTTWVQCPAPDAAAE PAGAARWAVIGAEVPAFAEAVRRRGGVAETFPDPAAARAAGPFDGHLLTAPAPDPDAPAEAAAESPLRTAEGPPAARTLLLTRSGVS TGPDDTPDPTASAVRARLLARGGDGTVLVDLGPEDPRPTGGLAEAEAEAEAEAEADGAFAGAAARDADTGDAPGSDVPSPDVEAWLA VLSAPAGETEFALRSGAALVPRLARVRAAGEVALPSGGPGAALVTGAGTGISRVLATHLVVNHGVRDLVLADAGEEGDPGAVATLAA DLTGLGATVTVRTGRPDTAERASALLGALGPDRPFALVVHPVTAPADAEAAGRLLDLTRRSEGTTLVLCGRAAPGDPSEAAAAAHAA ALTHDVAHATFLAFGPVAPDAPQPGLAALTAREAGDALDAVRATAGGPALVLRTDPWTLRAPGAERLRRMLPRYAEDTAAPAALDEG LRERLGVLPGRARQRALTALVREEAAGVLGLDAPRRIDAGLAFTRLGLTSLTAVALRDRLAARTGLRLPVTLAFDHPTPAAVAAVLD GELFGARPATDAARRTSEDAGTSHRTGPRHDAHDPVVVIGMSCRYPGGIASPDDLWRFVAEGRDAIGPFPTDRGWDLERLLSGDGTA PGTSATGSGGFLDAPGDFDAAFFGVSPREALAMDPQQRLLLEVGWEAVERAGIDPTSLRHTDTGVYVGLMFHDYAQHTGDLPEDLER LLGTGTAGSVASGRLAYTLGLNGPALTVDTACSSSLVAVHLAAQALRRGDCSLALAGGVAVMATPSTFVEFSRQHGLAADGRCKAFG EGADGTGWSEGVGVVALERLSDARRHGHPVLAVIRGSAVNQDGASNGLTAPSGPAQQQVIRRALAAAGLTPGDVDAVEAHGTGTVLG DPIEANALLATYGRGRDPERPLLLGSFKSNIGHAQSAAGVGAVIKMVQAIRYGELPRTLHAERPTPAVDWSGGAVRLLREHTPWPVT GDRRRAAVSSFGVSGTNVHLVLEQAPEPDAPPALPPAVHPDTMPVLLSARTPSALRRQAERVLDAVRADPGLPVPDLAAALATTRTA FPRRAAFVVEGRAELERRLRSFADGEDGGAEPDEVPAEPSLAIGFTGQGSQHPGMGRELYAAFPVFAEALDAAWAALDPHLERPLRD VMWAPDGTARAALLDRTDFTQAALFALEGALYRLVESWGVVPDVVLGHSVGALTAAHAAGVLSLPDAASLVAARGRLMAALPPGGAM TSIEATEDELRTELRATDGVLAIAAVNGPRSVVVSGEEKAVRTVGEAFRARGRRVTALRVSHAFHSPLMDPMVEEFRAAAARVTYRP PLLPMISDLTGRPADPAHLRSPDHWVRHVRETVRFADAVRALPGQGVTAFLELGPDRQLTTMAAAGAPGSGPALCGGLRRGRSEVRS LLDAVAQAHVRGVPVDWREFFAGGPTRPVDLPVYPFEHRRYWLASAPTRTGGATSVPEAPAPPTGDTPGPDRLRADLAPLTDDERER RLTDLVRGEIAAVAGFDGPHEIEAHRSMTDLGLDSVSAVDLSTRLGARTGLDLPASLAFDHPTSTAIARHLLAALRTQGLTPESGQD AGTDGGTEGGKHAGIRAAEGTVFGALARLEASVDRDRPDAAARDRLVERLRALTARLTEGPAGGPSADDTSIGDRIGEATTVDQLFD LIDNELKGPLEHHHHHHHH |

| Construct | Amino acid sequence |
|---|---|
| VemG(M0-M1)<br>JH015 | MSAWSHPQFEKGGGSGGGSGGSAWSHPQFEKGAGSVTQVDDIRPLSAVLRAHALDRPDKVAFADEERHVTYARLAERTGRLAGHLAG<br>HGLRRGDRVAILLGNSVTTVESYLAVTRAAAVGVPVNPQSSDAELAHQLDDSGARFVITDGPHLDQVERLRATREGIRVVLARGYGP<br>GGGPHGPGPEAPASEGAPALDGTSAPDGAPGPDGPLSFEELAGSEPAQAPRDDLGLDEPAWILYTSGTTGAAKGVVSHQRACLWSVR<br>SSYQGVLGLGPDDRLLWPLPLFHSLAHILCVLGVTVSGATARILPGFGARDVLDALRAEPCTMLLGVPSMYLRLVAAVEAVEEAGGS<br>VPKPRLCVVTGAATGPELASAVERVLGAPLVNSYGATETCGAITLSRPDSERPPGTVGTAVPGSELRLVDPRTGRDARTGDEGEVLV<br>RSPGLLLRYHDRPEETAAALRDGWYRTGDLARRDAAGRLTITGRVKELVIRAGENIQPGEIEDVLRTVPGIADAAVTGRPDEALGEV<br>PVAYVVPAAGGWSPAEALAACRERLSYFKVPAELYEIAALPVTGSGKVTRRALPGLPARLRALGTGHHERLWRTTWVQCPAPDAAAE<br>PAGAARWAVIGAEVPAFAEAVRRRGGVAETFPDPAAARAAGPFDGHLLTAPAPDPDAPAEAAAESPLRTAEGPPAARTLLLTRSGVS<br>TGPDDTPDPTASAVRARLLARGGDGTVLVDLGPEDPRPTGGLAEAEAEAEAEAEADGAFAGAAARDADTGDAPGSDVPSPDVEAWLA<br>VLSAPAGETEFALRSGAALVPRLARVRAAGEVALPSGGPGAALVTGAGTGISRVLATHLVVNHGVRDLVLADAGEEGDPGAVATLAA<br>DLTGLGATVTVRTGRPDTAERASALLGALGPDRPFALVVHPVTAPADAEAAGRLLDLTRRSEGTTLVLCGRAAPGDPSEAAAAAHAA<br>ALTHDVAHATFLAFGPVAPDAPQPGLAALTAREAGDALDAVRATAGGPALVLRTDPWTLRAPGAERLRRMLPRYAEDTAAPAALDEG<br>LRERLGVLPGRARQRALTALVREEAAGVLGLDAPRRIDAGLAFTRLGLTSLTAVALRDRLAARTGLRLPVTLAFDHPTPAAVAAVLD<br>GELFGARPATDAARRTSEDAGTSHRTGPRHDAHDPVVVIGMSCRYPGGIASPDDLWRFVAEGRDAIGPFPTDRGWDLERLLSGDGTA<br>PGTSATGSGGFLDAPGDFDAAFFGVSPREALAMDPQQRLLLEVGWEAVERAGIDPTSLRHTDTGVYVGLMFHDYAQHTGDLPEDLER<br>LLGTGTAGSVASGRLAYTLGLNGPALTVDTACSSSLVAVHLAAQALRRGDCSLALAGGVAVMATPSTFVEFSRQHGLAADGRCKAFG<br>EGADGTGWSEGVGVVALERLSDARRHGHPVLAVIRGSAVNQDGASNGLTAPSGPAQQQVIRRALAAAGLTPGDVDAVEAHGTGTVLG<br>DPIEANALLATYGRGRDPERPLLLGSFKSNIGHAQSAAGVGAVIKMVQAIRYGELPRTLHAERPTPAVDWSGGAVRLLREHTPWPVT<br>GDRRRAAVSSFGVSGTNVHLVLEQAPEPDAPPALPPAVHPDTMPVLLSARTPSALRRQAERVLDAVRADPGLPVPDLAAALATTRTA<br>FPRRAAFVVEGRAELERRLRSFADGEDGGAEPDEVPAEPSLAIGFTGQGSQHPGMGRELYAAFPVFAEALDAAWAALDPHLERPLRD<br>VMWAPDGTARAALLDRTDFTQAALFALEGALYRLVESWGVVPDVVLGHSVGALTAAHAAGVLSLPDAASLVAARGRLMAALPPGGAM<br>TSIEATEDELRTELRATDGVLAIAAVNGPRSVVVSGEEKAVRTVGEAFRARGRRVTALRVSHAFHSPLMDPMVEEFRAAAARVTYRP<br>PLLPMISDLTGRPADPAHLRSPDHWVRHVRETVRFADAVRALPGQGVTAFLELGPDRQLTTMAAAGAPGSGPALCGGLRRGRSEVRS<br>LLDAVAQAHVRGVPVDWREFFAGGPTRPVDLPVYPFEHRRYWLASAPTRTGGATSVPEAPAPPTGDTPGPDRLRADLAPLTDDERER<br>RLTDLVRGEIAAVAGFDGPHEIEAHRSMTDLGLDSVSAVDLSTRLGARTGLDLPASLAFDHPTSTAIARHLLAALRLEHHHHHHHH |
| VemG(M0-M1)-DEBS(DD2)<br>JH013 | MSAWSHPQFEKGGGSGGGSGGSAWSHPQFEKGAGSVTQVDDIRPLSAVLRAHALDRPDKVAFADEERHVTYARLAERTGRLAGHLAG<br>HGLRRGDRVAILLGNSVTTVESYLAVTRAAAVGVPVNPQSSDAELAHQLDDSGARFVITDGPHLDQVERLRATREGIRVVLARGYGP<br>GGGPHGPGPEAPASEGAPALDGTSAPDGAPGPDGPLSFEELAGSEPAQAPRDDLGLDEPAWILYTSGTTGAAKGVVSHQRACLWSVR<br>SSYQGVLGLGPDDRLLWPLPLFHSLAHILCVLGVTVSGATARILPGFGARDVLDALRAEPCTMLLGVPSMYLRLVAAVEAVEEAGGS<br>VPKPRLCVVTGAATGPELASAVERVLGAPLVNSYGATETCGAITLSRPDSERPPGTVGTAVPGSELRLVDPRTGRDARTGDEGEVLV<br>RSPGLLLRYHDRPEETAAALRDGWYRTGDLARRDAAGRLTITGRVKELVIRAGENIQPGEIEDVLRTVPGIADAAVTGRPDEALGEV<br>PVAYVVPAAGGWSPAEALAACRERLSYFKVPAELYEIAALPVTGSGKVTRRALPGLPARLRALGTGHHERLWRTTWVQCPAPDAAAE<br>PAGAARWAVIGAEVPAFAEAVRRRGGVAETFPDPAAARAAGPFDGHLLTAPAPDPDAPAEAAAESPLRTAEGPPAARTLLLTRSGVS<br>TGPDDTPDPTASAVRARLLARGGDGTVLVDLGPEDPRPTGGLAEAEAEAEAEAEADGAFAGAAARDADTGDAPGSDVPSPDVEAWLA<br>VLSAPAGETEFALRSGAALVPRLARVRAAGEVALPSGGPGAALVTGAGTGISRVLATHLVVNHGVRDLVLADAGEEGDPGAVATLAA<br>DLTGLGATVTVRTGRPDTAERASALLGALGPDRPFALVVHPVTAPADAEAAGRLLDLTRRSEGTTLVLCGRAAPGDPSEAAAAAHAA<br>ALTHDVAHATFLAFGPVAPDAPQPGLAALTAREAGDALDAVRATAGGPALVLRTDPWTLRAPGAERLRRMLPRYAEDTAAPAALDEG<br>LRERLGVLPGRARQRALTALVREEAAGVLGLDAPRRIDAGLAFTRLGLTSLTAVALRDRLAARTGLRLPVTLAFDHPTPAAVAAVLD<br>GELFGARPATDAARRTSEDAGTSHRTGPRHDAHDPVVVIGMSCRYPGGIASPDDLWRFVAEGRDAIGPFPTDRGWDLERLLSGDGTA<br>PGTSATGSGGFLDAPGDFDAAFFGVSPREALAMDPQQRLLLEVGWEAVERAGIDPTSLRHTDTGVYVGLMFHDYAQHTGDLPEDLER<br>LLGTGTAGSVASGRLAYTLGLNGPALTVDTACSSSLVAVHLAAQALRRGDCSLALAGGVAVMATPSTFVEFSRQHGLAADGRCKAFG<br>EGADGTGWSEGVGVVALERLSDARRHGHPVLAVIRGSAVNQDGASNGLTAPSGPAQQQVIRRALAAAGLTPGDVDAVEAHGTGTVLG<br>DPIEANALLATYGRGRDPERPLLLGSFKSNIGHAQSAAGVGAVIKMVQAIRYGELPRTLHAERPTPAVDWSGGAVRLLREHTPWPVT<br>GDRRRAAVSSFGVSGTNVHLVLEQAPEPDAPPALPPAVHPDTMPVLLSARTPSALRRQAERVLDAVRADPGLPVPDLAAALATTRTA<br>FPRRAAFVVEGRAELERRLRSFADGEDGGAEPDEVPAEPSLAIGFTGQGSQHPGMGRELYAAFPVFAEALDAAWAALDPHLERPLRD<br>VMWAPDGTARAALLDRTDFTQAALFALEGALYRLVESWGVVPDVVLGHSVGALTAAHAAGVLSLPDAASLVAARGRLMAALPPGGAM<br>TSIEATEDELRTELRATDGVLAIAAVNGPRSVVVSGEEKAVRTVGEAFRARGRRVTALRVSHAFHSPLMDPMVEEFRAAAARVTYRP<br>PLLPMISDLTGRPADPAHLRSPDHWVRHVRETVRFADAVRALPGQGVTAFLELGPDRQLTTMAAAGAPGSGPALCGGLRRGRSEVRS<br>LLDAVAQAHVRGVPVDWREFFAGGPTRPVDLPVYPFEHRRYWLASAPTRTGGATSVPEAPAPPTGDTPGPDRLRADLAPLTDDERER<br>RLTDLVRGEIAAVAGFDGPHEIEAHRSMTDLGLDSVSAVDLSTRLGARTGLDLPASLAFDHPTSTAIARHLLAALRTQGLTPESGQD<br>AGTDGGTEGGKHAGIRAAEGSALAGLDALEAALPEVPATEREELVQRLERMLAALRPVAQAADASGTGANPSGDDLGEAGVDELLEA<br>LGRELDGDLEHHHHHHHH |

| Construct | Amino acid sequence |
| --- | --- |
| VemG(M0-M1-DDa1-a2)-DEBS(a3)<br>JH014 | MSAWSHPQFEKGGGSGGGSGGSAWSHPQFEKGAGSVTQVDDIRPLSAVLRAHALDRPDKVAFADEERHVTYARLAERTGRLAGHLAG<br>HGLRRGDRVAILLGNSVTTVESYLAVTRAAAVGVPVNPQSSDAELAHQLDDSGARFVITDGPHLDQVERLRATREGIRVVLARGYGP<br>GGGPHGPGPEAPASEGAPALDGTSAPDGAPGPDGPLSFEELAGSEPAQAPRDDLGLDEPAWILYTSGTTGAAKGVVSHQRACLWSVR<br>SSYQGVLGLGPDDRLLWPLPLFHSLAHILCVLGVTVSGATARILPGFGARDVLDALRAEPCTMLLGVPSMYLRLVAAVEAVEEAGGS<br>VPKPRLCVVTGAATGPELASAVERVLGAPLVNSYGATETCGAITLSRPDSERPPGTVGTAVPGSELRLVDPRTGRDARTGDEGEVLV<br>RSPGLLLRYHDRPEETAAALRDGWYRTGDLARRDAAGRLTITGRVKELVIRAGENIQPGEIEDVLRTVPGIADAAVTGRPDEALGEV<br>PVAYVVPAAGGWSPAEALAACRERLSYFKVPAELYEIAALPVTGSGKVTRRALPGLPARLRALGTGHHERLWRTTWVQCPAPDAAAE<br>PAGAARWAVIGAEVPAFAEAVRRRGGVAETFPDPAAARAAGPFDGHLLTAPAPDPDAPAEAAAESPLRTAEGPPAARTLLLTRSGVS<br>TGPDDTPDPTASAVRARLLARGGDGTVLVDLGPEDPRPTGGLAEAEAEAEAEAEADGAFAGAAARDADTGDAPGSDVPSPDVEAWLA<br>VLSAPAGETEFALRSGAALVPRLARVRAAGEVALPSGGPGAALVTGAGTGISRVLATHLVVNHGVRDLVLADAGEEGDPGAVATLAA<br>DLTGLGATVTVRTGRPDTAERASALLGALGPDRPFALVVHPVTAPADAEAAGRLLDLTRRSEGTTLVLCGRAAPGDPSEAAAAAHAA<br>ALTHDVAHATFLAFGPVAPDAPQPGLAALTAREAGDALDAVRATAGGPALVLRTDPWTLRAPGAERLRRMLPRYAEDTAAPAALDEG<br>LRERLGVLPGRARQRALTALVREEAAGVLGLDAPRRIDAGLAFTRLGLTSLTAVALRDRLAARTGLRLPVTLAFDHPTPAAVAAVLD<br>GELFGARPATDAARRTSEDAGTSHRTGPRHDAHDPVVVIGMSCRYPGGIASPDDLWRFVAEGRDAIGPFPTDRGWDLERLLSGDGTA<br>PGTSATGSGGFLDAPGDFDAAFFGVSPREALAMDPQQRLLLEVGWEAVERAGIDPTSLRHTDTGVYVGLMFHDYAQHTGDLPEDLER<br>LLGTGTAGSVASGRLAYTLGLNGPALTVDTACSSSLVAVHLAAQALRRGDCSLALAGGVAVMATPSTFVEFSRQHGLAADGRCKAFG<br>EGADGTGWSEGVGVVALERLSDARRHGHPVLAVIRGSAVNQDGASNGLTAPSGPAQQQVIRRALAAAGLTPGDVDAVEAHGTGTVLG<br>DPIEANALLATYGRGRDPERPLLLGSFKSNIGHAQSAAGVGAVIKMVQAIRYGELPRTLHAERPTPAVDWSGGAVRLLREHTPWPVT<br>GDRRRAAVSSFGVSGTNVHLVLEQAPEPDAPPALPPAVHPDTMPVLLSARTPSALRRQAERVLDAVRADPGLPVPDLAAALATTRTA<br>FPRRAAFVVEGRAELERRLRSFADGEDGGAEPDEVPAEPSLAIGFTGQGSQHPGMGRELYAAFPVFAEALDAAWAALDPHLERPLRD<br>VMWAPDGTARAALLDRTDFTQAALFALEGALYRLVESWGVVPDVVLGHSVGALTAAHAAGVLSLPDAASLVAARGRLMAALPPGGAM<br>TSIEATEDELRTELRATDGVLAIAAVNGPRSVVVSGEEKAVRTVGEAFRARGRRVTALRVSHAFHSPLMDPMVEEFRAAAARVTYRP<br>PLLPMISDLTGRPADPAHLRSPDHWVRHVRETVRFADAVRALPGQGVTAFLELGPDRQLTTMAAAGAPGSGPALCGGLRRGRSEVRS<br>LLDAVAQAHVRGVPVDWREFFAGGPTRPVDLPVYPFEHRRYWLASAPTRTGGATSVPEAPAPPTGDTPGPDRLRADLAPLTDDERER<br>RLTDLVRGEIAAVAGFDGPHEIEAHRSMTDLGLDSVSAVDLSTRLGARTGLDLPASLAFDHPTSTAIARHLLAALRTQGLTPESGQD<br>AGTDGGTEGGKHAGIRAAEGTVFGALARLEASVDRDRPDAAARDRLVERLRALTARLTEGPAGADASGTGANPSGDDLGEAGVDELL<br>EALGRELDGDLEHHHHHHHH |
| VemG(M0-M1)-SZ3<br>CS002.2 | MSAWSHPQFEKGGGSGGGSGGSAWSHPQFEKGAGSVTQVDDIRPLSAVLRAHALDRPDKVAFADEERHVTYARLAERTGRLAGHLAG<br>HGLRRGDRVAILLGNSVTTVESYLAVTRAAAVGVPVNPQSSDAELAHQLDDSGARFVITDGPHLDQVERLRATREGIRVVLARGYGP<br>GGGPHGPGPEAPASEGAPALDGTSAPDGAPGPDGPLSFEELAGSEPAQAPRDDLGLDEPAWILYTSGTTGAAKGVVSHQRACLWSVR<br>SSYQGVLGLGPDDRLLWPLPLFHSLAHILCVLGVTVSGATARILPGFGARDVLDALRAEPCTMLLGVPSMYLRLVAAVEAVEEAGGS<br>VPKPRLCVVTGAATGPELASAVERVLGAPLVNSYGATETCGAITLSRPDSERPPGTVGTAVPGSELRLVDPRTGRDARTGDEGEVLV<br>RSPGLLLRYHDRPEETAAALRDGWYRTGDLARRDAAGRLTITGRVKELVIRAGENIQPGEIEDVLRTVPGIADAAVTGRPDEALGEV<br>PVAYVVPAAGGWSPAEALAACRERLSYFKVPAELYEIAALPVTGSGKVTRRALPGLPARLRALGTGHHERLWRTTWVQCPAPDAAAE<br>PAGAARWAVIGAEVPAFAEAVRRRGGVAETFPDPAAARAAGPFDGHLLTAPAPDPDAPAEAAAESPLRTAEGPPAARTLLLTRSGVS<br>TGPDDTPDPTASAVRARLLARGGDGTVLVDLGPEDPRPTGGLAEAEAEAEAEAEADGAFAGAAARDADTGDAPGSDVPSPDVEAWLA<br>VLSAPAGETEFALRSGAALVPRLARVRAAGEVALPSGGPGAALVTGAGTGISRVLATHLVVNHGVRDLVLADAGEEGDPGAVATLAA<br>DLTGLGATVTVRTGRPDTAERASALLGALGPDRPFALVVHPVTAPADAEAAGRLLDLTRRSEGTTLVLCGRAAPGDPSEAAAAAHAA<br>ALTHDVAHATFLAFGPVAPDAPQPGLAALTAREAGDALDAVRATAGGPALVLRTDPWTLRAPGAERLRRMLPRYAEDTAAPAALDEG<br>LRERLGVLPGRARQRALTALVREEAAGVLGLDAPRRIDAGLAFTRLGLTSLTAVALRDRLAARTGLRLPVTLAFDHPTPAAVAAVLD<br>GELFGARPATDAARRTSEDAGTSHRTGPRHDAHDPVVVIGMSCRYPGGIASPDDLWRFVAEGRDAIGPFPTDRGWDLERLLSGDGTA<br>PGTSATGSGGFLDAPGDFDAAFFGVSPREALAMDPQQRLLLEVGWEAVERAGIDPTSLRHTDTGVYVGLMFHDYAQHTGDLPEDLER<br>LLGTGTAGSVASGRLAYTLGLNGPALTVDTACSSSLVAVHLAAQALRRGDCSLALAGGVAVMATPSTFVEFSRQHGLAADGRCKAFG<br>EGADGTGWSEGVGVVALERLSDARRHGHPVLAVIRGSAVNQDGASNGLTAPSGPAQQQVIRRALAAAGLTPGDVDAVEAHGTGTVLG<br>DPIEANALLATYGRGRDPERPLLLGSFKSNIGHAQSAAGVGAVIKMVQAIRYGELPRTLHAERPTPAVDWSGGAVRLLREHTPWPVT<br>GDRRRAAVSSFGVSGTNVHLVLEQAPEPDAPPALPPAVHPDTMPVLLSARTPSALRRQAERVLDAVRADPGLPVPDLAAALATTRTA<br>FPRRAAFVVEGRAELERRLRSFADGEDGGAEPDEVPAEPSLAIGFTGQGSQHPGMGRELYAAFPVFAEALDAAWAALDPHLERPLRD<br>VMWAPDGTARAALLDRTDFTQAALFALEGALYRLVESWGVVPDVVLGHSVGALTAAHAAGVLSLPDAASLVAARGRLMAALPPGGAM<br>TSIEATEDELRTELRATDGVLAIAAVNGPRSVVVSGEEKAVRTVGEAFRARGRRVTALRVSHAFHSPLMDPMVEEFRAAAARVTYRP<br>PLLPMISDLTGRPADPAHLRSPDHWVRHVRETVRFADAVRALPGQGVTAFLELGPDRQLTTMAAAGAPGSGPALCGGLRRGRSEVRS<br>LLDAVAQAHVRGVPVDWREFFAGGPTRPVDLPVYPFEHRRYWLASAPTRTGGATSVPEAPAPPTGDTPGPDRLRADLAPLTDDERER<br>RLTDLVRGEIAAVAGFDGPHEIEAHRSMTDLGLDSVSAVDLSTRLGARTGLDLPASLAFDHPTSTAIARHLLAALGGSGGGSGNEVT<br>TLENDAAFIENENAYLEKEIARLRKEKAALRNRLAHKKLEHHHHHH |

| Construct | Amino acid sequence |
|---|---|
| VemG(M0-M1-a1-a2)-SpyC<br>JH017 | MSAWSHPQFEKGGGSGGGSGGSAWSHPQFEKGAGSVTQVDDIRPLSAVLRAHALDRPDKVAFADEERHVTYARLAERTGRLAGHLAG<br>HGLRRGDRVAILLGNSVTTVESYLAVTRAAAVGVPVNPQSSDAELAHQLDDSGARFVITDGPHLDQVERLRATREGIRVVLARGYGP<br>GGGPHGPGPEAPASEGAPALDGTSAPDGAPGPDGPLSFEELAGSEPAQAPRDDLGLDEPAWILYTSGTTGAAKGVVSHQRACLWSVR<br>SSYQGVLGLGPDDRLLWPLPLFHSLAHILCVLGVTVSGATARILPGFGARDVLDALRAEPCTMLLGVPSMYLRLVAAVEAVEEAGGS<br>VPKPRLCVVTGAATGPELASAVERVLGAPLVNSYGATETCGAITLSRPDSERPPGTVGTAVPGSELRLVDPRTGRDARTGDEGEVLV<br>RSPGLLLRYHDRPEETAAALRDGWYRTGDLARRDAAGRLTITGRVKELVIRAGENIQPGEIEDVLRTVPGIADAAVTGRPDEALGEV<br>PVAYVVPAAGGWSPAEALAACRERLSYFKVPAELYEIAALPVTGSGKVTRRALPGLPARLRALGTGHHERLWRTTWVQCPAPDAAAE<br>PAGAARWAVIGAEVPAFAEAVRRRGGVAETFPDPAAARAAGPFDGHLLTAPAPDPDAPAEAAAESPLRTAEGPPAARTLLLTRSGVS<br>TGPDDTPDPTASAVRARLLARGGDGTVLVDLGPEDPRPTGGLAEAEAEAEAEAEADGAFAGAAARDADTGDAPGSDVPSPDVEAWLA<br>VLSAPAGETEFALRSGAALVPRLARVRAAGEVALPSGGPGAALVTGAGTGISRVLATHLVVNHGVRDLVLADAGEEGDPGAVATLAA<br>DLTGLGATVTVRTGRPDTAERASALLGALGPDRPFALVVHPVTAPADAEAAGRLLDLTRRSEGTTLVLCGRAAPGDPSEAAAAAHAA<br>ALTHDVAHATFLAFGPVAPDAPQPGLAALTAREAGDALDAVRATAGGPALVLRTDPWTLRAPGAERLRRMLPRYAEDTAAPAALDEG<br>LRERLGVLPGRARQRALTALVREEAAGVLGLDAPRRIDAGLAFTRLGLTSLTAVALRDRLAARTGLRLPVTLAFDHPTPAAVAAVLD<br>GELFGARPATDAARRTSEDAGTSHRTGPRHDAHDPVVVIGMSCRYPGGIASPDDLWRFVAEGRDAIGPFPTDRGWDLERLLSGDGTA<br>PGTSATGSGGFLDAPGDFDAAFFGVSPREALAMDPQQRLLLEVGWEAVERAGIDPTSLRHTDTGVYVGLMFHDYAQHTGDLPEDLER<br>LLGTGTAGSVASGRLAYTLGLNGPALTVDTACSSSLVAVHLAAQALRRGDCSLALAGGVAVMATPSTFVEFSRQHGLAADGRCKAFG<br>EGADGTGWSEGVGVVALERLSDARRHGHPVLAVIRGSAVNQDGASNGLTAPSGPAQQQVIRRALAAAGLTPGDVDAVEAHGTGTVLG<br>DPIEANALLATYGRGRDPERPLLLGSFKSNIGHAQSAAGVGAVIKMVQAIRYGELPRTLHAERPTPAVDWSGGAVRLLREHTPWPVT<br>GDRRRAAVSSFGVSGTNVHLVLEQAPEPDAPPALPPAVHPDTMPVLLSARTPSALRRQAERVLDAVRADPGLPVPDLAAALATTRTA<br>FPRRAAFVVEGRAELERRLRSFADGEDGGAEPDEVPAEPSLAIGFTGQGSQHPGMGRELYAAFPVFAEALDAAWAALDPHLERPLRD<br>VMWAPDGTARAALLDRTDFTQAALFALEGALYRLVESWGVVPDVVLGHSVGALTAAHAAGVLSLPDAASLVAARGRLMAALPPGGAM<br>TSIEATEDELRTELRATDGVLAIAAVNGPRSVVVSGEEKAVRTVGEAFRARGRRVTALRVSHAFHSPLMDPMVEEFRAAAARVTYRP<br>PLLPMISDLTGRPADPAHLRSPDHWVRHVRETVRFADAVRALPGQGVTAFLELGPDRQLTTMAAAGAPGSGPALCGGLRRGRSEVRS<br>LLDAVAQAHVRGVPVDWREFFAGGPTRPVDLPVYPFEHRRYWLASAPTRTGGATSVPEAPAPPTGDTPGPDRLRADLAPLTDDERER<br>RLTDLVRGEIAAVAGFDGPHEIEAHRSMTDLGLDSVSAVDLSTRLGARTGLDLPASLAFDHPTSTAIARHLLAALRTQGLTPESGQD<br>AGTDGGTEGGKHAGIRAAEGTVFGALARLEASVDRDRPDAAARDRLVERLRALTARLTEGPAGGPSADDTSIGGGSGAMVDTLSGLS<br>SEQGQSGDMTIEEDSATHIKFSKRDEDGKELAGATMELRDSSGKTISTWISDGQVKDFYLYPGKYTFVETAAPDGYEVATAITFTVN<br>EQGQVTVNGKATKGDAHILEHHHHHHHH |
| VemG(M0)-DEBS(DD4)<br>LB047 | MSAWSHPQFEKGGGSGGGSGGSAWSHPQFEKGAGSVTQVDDIRPLSAVLRAHALDRPDKVAFADEERHVTYARLAERTGRLAGHLAG<br>HGLRRGDRVAILLGNSVTTVESYLAVTRAAAVGVPVNPQSSDAELAHQLDDSGARFVITDGPHLDQVERLRATREGIRVVLARGYGP<br>GGGPHGPGPEAPASEGAPALDGTSAPDGAPGPDGPLSFEELAGSEPAQAPRDDLGLDEPAWILYTSGTTGAAKGVVSHQRACLWSVR<br>SSYQGVLGLGPDDRLLWPLPLFHSLAHILCVLGVTVSGATARILPGFGARDVLDALRAEPCTMLLGVPSMYLRLVAAVEAVEEAGGS<br>VPKPRLCVVTGAATGPELASAVERVLGAPLVNSYGATETCGAITLSRPDSERPPGTVGTAVPGSELRLVDPRTGRDARTGDEGEVLV<br>RSPGLLLRYHDRPEETAAALRDGWYRTGDLARRDAAGRLTITGRVKELVIRAGENIQPGEIEDVLRTVPGIADAAVTGRPDEALGEV<br>PVAYVVPAAGGWSPAEALAACRERLSYFKVPAELYEIAALPVTGSGKVTRRALPGLPARLRALGTGHHERLWRTTWVQCPAPDAAAE<br>PAGAARWAVIGAEVPAFAEAVRRRGGVAETFPDPAAARAAGPFDGHLLTAPAPDPDAPAEAAAESPLRTAEGPPAARTLLLTRSGVS<br>TGPDDTPDPTASAVRARLLARGGDGTVLVDLGPEDPRPTGGLAEAEAEAEAEAEADGAFAGAAARDADTGDAPGSDVPSPDVEAWLA<br>VLSAPAGETEFALRSGAALVPRLARVRAAGEVALPSGGPGAALVTGAGTGISRVLATHLVVNHGVRDLVLADAGEEGDPGAVATLAA<br>DLTGLGATVTVRTGRPDTAERASALLGALGPDRPFALVVHPVTAPADAEAAGRLLDLTRRSEGTTLVLCGRAAPGDPSEAAAAAHAA<br>ALTHDVAHATFLAFGPVAPDAPQPGLAALTAREAGDALDAVRATAGGPALVLRTDPWTLRAPGAERLRRMLPRYAEDTAAPAALDEG<br>LRERLGVLPGRARQRALTALVREEAAGVLGLDAPRRIDAGLAFTRLGLTSLTAVALRDRLAARTGLRLPVTLAFDHPTPAAVAAVLD<br>GELFAASPAVDIGDRLDELEKALEALSAEDGHDDVGQRLESLLRRWNSRRADAPSTSAISEDASDDELFSMLDQRFGGGEDLPNSSS<br>VDKLAAALEHHHHHH |
| DEBS(DD5)-VemG(M1-DD1)<br>CS008 | MSGDNGMTEEKLRRYLKRTVTELDSVTARLREVEHRAGDPVVVIGMSCRYPGGIASPDDLWRFVAEGRDAIGPFPTDRGWDLERLLS<br>GDGTAPGTSATGSGGFLDAPGDFDAAFFGVSPREALAMDPQQRLLLEVGWEAVERAGIDPTSLRHTDTGVYVGLMFHDYAQHTGDLP<br>EDLERLLGTGTAGSVASGRLAYTLGLNGPALTVDTACSSSLVAVHLAAQALRRGDCSLALAGGVAVMATPSTFVEFSRQHGLAADGR<br>CKAFGEGADGTGWSEGVGVVALERLSDARRHGHPVLAVIRGSAVNQDGASNGLTAPSGPAQQQVIRRALAAAGLTPGDVDAVEAHGT<br>GTVLGDPIEANALLATYGRGRDPERPLLLGSFKSNIGHAQSAAGVGAVIKMVQAIRYGELPRTLHAERPTPAVDWSGGAVRLLREHT<br>PWPVTGDRRRAAVSSFGVSGTNVHLVLEQAPEPDAPPALPPAVHPDTMPVLLSARTPSALRRQAERVLDAVRADPGLPVPDLAAALA<br>TTRTAFPRRAAFVVEGRAELERRLRSFADGEDGGAEPDEVPAEPSLAIGFTGQGSQHPGMGRELYAAFPVFAEALDAAWAALDPHLE<br>RPLRDVMWAPDGTARAALLDRTDFTQAALFALEGALYRLVESWGVVPDVVLGHSVGALTAAHAAGVLSLPDAASLVAARGRLMAALP<br>PGGAMTSIEATEDELRTELRATDGVLAIAAVNGPRSVVVSGEEKAVRTVGEAFRARGRRVTALRVSHAFHSPLMDPMVEEFRAAAAR<br>VTYRPPLLPMISDLTGRPADPAHLRSPDHWVRHVRETVRFADAVRALPGQGVTAFLELGPDRQLTTMAAAGAPGSGPALCGGLRRGR<br>SEVRSLLDAVAQAHVRGVPVDWREFFAGGPTRPVDLPVYPFEHRRYWLASAPTRTGGATSVPEAPAPPTGDTPGPDRLRADLAPLTD<br>DERERRLTDLVRGEIAAVAGFDGPHEIEAHRSMTDLGLDSVSAVDLSTRLGARTGLDLPASLAFDHPTSTAIARHLLAALRTQGLTP<br>ESGQDAGTDGGTEGGKHAGIRAAEGTVFGALARLEASVDRDRPDAAARDRLVERLRALTARLTEGPAGGPSADDTSIGDRIGEATTV<br>DQLFDLIDNELKGPLEHHHHHH |
| DEBS(DD5)-VemG(M1)-SZ3<br>LB048 | MSGDNGMTEEKLRRYLKRTVTELDSVTARLREVEHRAGDPVVVIGMSCRYPGGIASPDDLWRFVAEGRDAIGPFPTDRGWDLERLLS<br>GDGTAPGTSATGSGGFLDAPGDFDAAFFGVSPREALAMDPQQRLLLEVGWEAVERAGIDPTSLRHTDTGVYVGLMFHDYAQHTGDLP<br>EDLERLLGTGTAGSVASGRLAYTLGLNGPALTVDTACSSSLVAVHLAAQALRRGDCSLALAGGVAVMATPSTFVEFSRQHGLAADGR<br>CKAFGEGADGTGWSEGVGVVALERLSDARRHGHPVLAVIRGSAVNQDGASNGLTAPSGPAQQQVIRRALAAAGLTPGDVDAVEAHGT<br>GTVLGDPIEANALLATYGRGRDPERPLLLGSFKSNIGHAQSAAGVGAVIKMVQAIRYGELPRTLHAERPTPAVDWSGGAVRLLREHT<br>PWPVTGDRRRAAVSSFGVSGTNVHLVLEQAPEPDAPPALPPAVHPDTMPVLLSARTPSALRRQAERVLDAVRADPGLPVPDLAAALA<br>TTRTAFPRRAAFVVEGRAELERRLRSFADGEDGGAEPDEVPAEPSLAIGFTGQGSQHPGMGRELYAAFPVFAEALDAAWAALDPHLE<br>RPLRDVMWAPDGTARAALLDRTDFTQAALFALEGALYRLVESWGVVPDVVLGHSVGALTAAHAAGVLSLPDAASLVAARGRLMAALP<br>PGGAMTSIEATEDELRTELRATDGVLAIAAVNGPRSVVVSGEEKAVRTVGEAFRARGRRVTALRVSHAFHSPLMDPMVEEFRAAAAR<br>VTYRPPLLPMISDLTGRPADPAHLRSPDHWVRHVRETVRFADAVRALPGQGVTAFLELGPDRQLTTMAAAGAPGSGPALCGGLRRGR<br>SEVRSLLDAVAQAHVRGVPVDWREFFAGGPTRPVDLPVYPFEHRRYWLASAPTRTGGATSVPEAPAPPTGDTPGPDRLRADLAPLTD<br>DERERRLTDLVRGEIAAVAGFDGPHEIEAHRSMTDLGLDSVSAVDLSTRLGARTGLDLPASLAFDHPTSTAIARHLLAALGGSGGGS |

| Construct | Amino acid sequence |
|---|---|
| | GNEVTTLENDAAFIENENAYLEKEIARLRKEKAALRNRLAHKKLEHHHHHH |

**Table S5.** Amino acid sequences of VemH and VemH-based constructs

| Construct | Amino acid sequence |
|---|---|
| VemH(DD2-M2-TE) CS005 | MTGTEEKLVDYLKKVTAELQETRRQLRGALAASREPIAIVGMACRYPGGVRTPEALWRLVLDEQDAISGFPTNRGWDIDGIYHPDPDRPGTCYAREGGFLHDAALFDAEFFGVSPREAQAMDPQQRLLLETAWEAFERAGIDPTSLRGSDTGVFAGVVHHDYATARVPETLEPYLVTGLSGGVASGRIAYTFGFEGPAVTVDTACSSSLVALHQAAHALRSGECELALAGGVTIMATPRAFLSFSRQRGLSPDGRCRAFGAGADGTGWAEGAGMLLVERLSDARRKGHPVLAVLRGSAVNQDGASNGLSAPNGPSQQRVIRKALAHAGLAARDVDVVEGHGTGTKLGDPIEAQALLATYGQERDAGLPLHLGSMKSNVGHSQAAAGVGGVIKMIEAMRHGILPRTLHADEPTPHVDWSAGDIELLTRRRAWPETGRPRRAAVSSFGISGTNAHVILEAPPEEPARDAAETRPEARSREAAEGRREAAGQGRTETGPGPSATPPEAPGRARPPVPWPLSGRDAGALRDQIGRLRAHLDAAPADPEDVAHSLARRAVFRHRAVLLAAPQAPAGGSPRAVTGVARPGGTALLFSGQGSQRVGMGSELYETYPVFAESFDAVAEHTGLPLKDVVLGGTPDGLLDRTRYAQPALFAVEVSLFRLVRALGLDVRAVVGHSVGEIAAAHVAGVMSMADACRLVEARGRLMDALPPGGAMVAVEVTEAEASAALAGLEDRVAVAAVNGPASTVLSGEEGAVLKLADAWRERGVRTHRLTVSHAFHSPLMEPMVDAFREVVAGLDLHRPTLAGLPAEVVDPEYWVRHVRRPVRFADAVARAREAGAVRWLEVGPGGVLTALAQRIVPDTEEHVFAAALRTDRPEPEALLVALSQVHVDGGTVDWSGLCAGGRLVDLPTYPFQRQHYWIEDQPLPPTAPRPGTAPSGTGTAAEGAAAAEVPLSERLARLTGAERLAAVRELVLAEASETLGHTGTLITADRTRQELGFDSLTAIELRNRISRTLGVRLPPTLVFDHEDLGEIASFVDARLDDAATGRSTGHGPLGEDGSGLLTELFREAAAAGRLDDAVTLTEAAARMRRTFTDAEDPAVRRTPVWFGRGPARPTVVCLPSFSAIAGVHVYARFADAFGDGWRVAALAHPGFVPGEPLPDSVDVLAELHARTVLDTVGADPFLLVGRSAGGWVAHEVAAVLERMGRAPDGVALLDTPARADDPRGHAVMVGGMLERDSRLVTIDDYRLTAMGGYSRLFREWKPEPIAAATLLVHAATPYGADEARIASWDLPHQAVKVTGDHFTMLERHSATTAEAVEQWSRSLLKLAAALEHHHHHHHH |
| VemH(M2-TE) JH008 | MEPIAIVGMACRYPGGVRTPEALWRLVLDEQDAISGFPTNRGWDIDGIYHPDPDRPGTCYAREGGFLHDAALFDAEFFGVSPREAQAMDPQQRLLLETAWEAFERAGIDPTSLRGSDTGVFAGVVHHDYATARVPETLEPYLVTGLSGGVASGRIAYTFGFEGPAVTVDTACSSSLVALHQAAHALRSGECELALAGGVTIMATPRAFLSFSRQRGLSPDGRCRAFGAGADGTGWAEGAGMLLVERLSDARRKGHPVLAVLRGSAVNQDGASNGLSAPNGPSQQRVIRKALAHAGLAARDVDVVEGHGTGTKLGDPIEAQALLATYGQERDAGLPLHLGSMKSNVGHSQAAAGVGGVIKMIEAMRHGILPRTLHADEPTPHVDWSAGDIELLTRRRAWPETGRPRRAAVSSFGISGTNAHVILEAPPEEPARDAAETRPEARSREAAEGRREAAGQGRTETGPGPSATPPEAPGRARPPVPWPLSGRDAGALRDQIGRLRAHLDAAPADPEDVAHSLARRAVFRHRAVLLAAPQAPAGGSPRAVTGVARPGGTALLFSGQGSQRVGMGSELYETYPVFAESFDAVAEHTGLPLKDVVLGGTPDGLLDRTRYAQPALFAVEVSLFRLVRALGLDVRAVVGHSVGEIAAAHVAGVMSMADACRLVEARGRLMDALPPGGAMVAVEVTEAEASAALAGLEDRVAVAAVNGPASTVLSGEEGAVLKLADAWRERGVRTHRLTVSHAFHSPLMEPMVDAFREVVAGLDLHRPTLAGLPAEVVDPEYWVRHVRRPVRFADAVARAREAGAVRWLEVGPGGVLTALAQRIVPDTEEHVFAAALRTDRPEPEALLVALSQVHVDGGTVDWSGLCAGGRLVDLPTYPFQRQHYWIEDQPLPPTAPRPGTAPSGTGTAAEGAAAAEVPLSERLARLTGAERLAAVRELVLAEASETLGHTGTLITADRTRQELGFDSLTAIELRNRISRTLGVRLPPTLVFDHEDLGEIASFVDARLDDAATGRSTGHGPLGEDGSGLLTELFREAAAAGRLDDAVTLTEAAARMRRTFTDAEDPAVRRTPVWFGRGPARPTVVCLPSFSAIAGVHVYARFADAFGDGWRVAALAHPGFVPGEPLPDSVDVLAELHARTVLDTVGADPFLLVGRSAGGWVAHEVAAVLERMGRAPDGVALLDTPARADDPRGHAVMVGGMLERDSRLVTIDDYRLTAMGGYSRLFREWKPEPIAAATLLVHAATPYGADEARIASWDLPHQAVKVTGDHFTMLERHSATTAEAVEQWSRSLLKLAAALEHHHHHHHH |
| DEBS(DD3)-VemH(M2-TE) JH005 | MTDSEKVAEYLRRATLDLRAARQRIRELESEPIAIVGMACRYPGGVRTPEALWRLVLDEQDAISGFPTNRGWDIDGIYHPDPDRPGTCYAREGGFLHDAALFDAEFFGVSPREAQAMDPQQRLLLETAWEAFERAGIDPTSLRGSDTGVFAGVVHHDYATARVPETLEPYLVTGLSGGVASGRIAYTFGFEGPAVTVDTACSSSLVALHQAAHALRSGECELALAGGVTIMATPRAFLSFSRQRGLSPDGRCRAFGAGADGTGWAEGAGMLLVERLSDARRKGHPVLAVLRGSAVNQDGASNGLSAPNGPSQQRVIRKALAHAGLAARDVDVVEGHGTGTKLGDPIEAQALLATYGQERDAGLPLHLGSMKSNVGHSQAAAGVGGVIKMIEAMRHGILPRTLHADEPTPHVDWSAGDIELLTRRRAWPETGRPRRAAVSSFGISGTNAHVILEAPPEEPARDAAETRPEARSREAAEGRREAAGQGRTETGPGPSATPPEAPGRARPPVPWPLSGRDAGALRDQIGRLRAHLDAAPADPEDVAHSLARRAVFRHRAVLLAAPQAPAGGSPRAVTGVARPGGTALLFSGQGSQRVGMGSELYETYPVFAESFDAVAEHTGLPLKDVVLGGTPDGLLDRTRYAQPALFAVEVSLFRLVRALGLDVRAVVGHSVGEIAAAHVAGVMSMADACRLVEARGRLMDALPPGGAMVAVEVTEAEASAALAGLEDRVAVAAVNGPASTVLSGEEGAVLKLADAWRERGVRTHRLTVSHAFHSPLMEPMVDAFREVVAGLDLHRPTLAGLPAEVVDPEYWVRHVRRPVRFADAVARAREAGAVRWLEVGPGGVLTALAQRIVPDTEEHVFAAALRTDRPEPEALLVALSQVHVDGGTVDWSGLCAGGRLVDLPTYPFQRQHYWIEDQPLPPTAPRPGTAPSGTGTAAEGAAAAEVPLSERLARLTGAERLAAVRELVLAEASETLGHTGTLITADRTRQELGFDSLTAIELRNRISRTLGVRLPPTLVFDHEDLGEIASFVDARLDDAATGRSTGHGPLGEDGSGLLTELFREAAAAGRLDDAVTLTEAAARMRRTFTDAEDPAVRRTPVWFGRGPARPTVVCLPSFSAIAGVHVYARFADAFGDGWRVAALAHPGFVPGEPLPDSVDVLAELHARTVLDTVGADPFLLVGRSAGGWVAHEVAAVLERMGRAPDGVALLDTPARADDPRGHAVMVGGMLERDSRLVTIDDYRLTAMGGYSRLFREWKPEPIAAATLLVHAATPYGADEARIASWDLPHQAVKVTGDHFTMLERHSATTAEAVEQWSRSLLKLAAALEHHHHHHHH |

| Construct | Amino acid sequence |
| --- | --- |
| SZ4-VemH(M2-TE) CS003 | MQKVAELKNRVAVKLNRNEQLKNKVEELKNRNAYLKNELATLENEVARLENDVAEGGSGGGSGKLEPIAIVGMACRYPGGVRTPEAL<br>WRLVLDEQDAISGFPTNRGWDIDGIYHPDPDRPGTCYAREGGFLHDAALFDAEFFGVSPREAQAMDPQQRLLLETAWEAFERAGIDP<br>TSLRGSDTGVFAGVVHHDYATARVPETLEPYLVTGLSGGVASGRIAYTFGFEGPAVTVDTACSSSLVALHQAAHALRSGECELALAG<br>GVTIMATPRAFLSFSRQRGLSPDGRCRAFGAGADGTGWAEGAGMLLVERLSDARRKGHPVLAVLRGSAVNQDGASNGLSAPNGPSQQ<br>RVIRKALAHAGLAARDVDVVEGHGTGTKLGDPIEAQALLATYGQERDAGLPLHLGSMKSNVGHSQAAAGVGGVIKMIEAMRHGILPR<br>TLHADEPTPHVDWSAGDIELLTRRRAWPETGRPRRAAVSSFGISGTNAHVILEAPPEEPARDAAETRPEARSREAAEGRREAAGQGR<br>TETGPGPSATPPEAPGRARPPVPWPLSGRDAGALRDQIGRLRAHLDAAPADPEDVAHSLARRAVFRHRAVLLAAPQAPAGGSPRAVT<br>GVARPGGTALLFSGQGSQRVGMGSELYETYPVFAESFDAVAEHTGLPLKDVVLGGTPDGLLDRTRYAQPALFAVEVSLFRLVRALGL<br>DVRAVVGHSVGEIAAAHVAGVMSMADACRLVEARGRLMDALPPGGAMVAVEVTEAEASAALAGLEDRVAVAAVNGPASTVLSGEEGA<br>VLKLADAWRERGVRTHRLTVSHAFHSPLMEPMVDAFREVVAGLDLHRPTLAGLPAEVVDPEYWVRHVRRPVRFADAVARAREAGAVR<br>WLEVGPGGVLTALAQRIVPDTEEHVFAAALRTDRPEPEALLVALSQVHVDGGTVDWSGLCAGGRLVDLPTYPFQRQHYWIEDQPLPP<br>TAPRPGTAPSGTGTAAEGAAAAEVPLSERLARLTGAERLAAVRELVLAEASETLGHTGTLITADRTRQELGFDSLTAIELRNRISRT<br>LGVRLPPTLVFDHEDLGEIASFVDARLDDAATGRSTGHGPLGEDGSGLLTELFREAAAAGRLDDAVTLTEAAARMRRTFTDAEDPAV<br>RRTPVWFGRGPARPTVVCLPSFSAIAGVHVYARFADAFGDGWRVAALAHPGFVPGEPLPDSVDVLAELHARTVLDTVGADPFLLVGR<br>SAGGWVAHEVAAVLERMGRAPDGVALLDTPARADDPRGHAVMVGGMLERDSRLVTIDDYRLTAMGGYSRLFREWKPEPIAAATLLVH<br>AATPYGADEARIASWDLPHQAVKVTGDHFTMLERHSATTAEAVEQWSRSLLKLAAALEHHHHHHHH |
| SpyT-VemH(M2-TE) JH010 | MAHIVMVDAYKPTKGGSGSPAAPAPASPASEPIAIVGMACRYPGGVRTPEALWRLVLDEQDAISGFPTNRGWDIDGIYHPDPDRPGT<br>CYAREGGFLHDAALFDAEFFGVSPREAQAMDPQQRLLLETAWEAFERAGIDPTSLRGSDTGVFAGVVHHDYATARVPETLEPYLVTG<br>LSGGVASGRIAYTFGFEGPAVTVDTACSSSLVALHQAAHALRSGECELALAGGVTIMATPRAFLSFSRQRGLSPDGRCRAFGAGADG<br>TGWAEGAGMLLVERLSDARRKGHPVLAVLRGSAVNQDGASNGLSAPNGPSQQRVIRKALAHAGLAARDVDVVEGHGTGTKLGDPIEA<br>QALLATYGQERDAGLPLHLGSMKSNVGHSQAAAGVGGVIKMIEAMRHGILPRTLHADEPTPHVDWSAGDIELLTRRRAWPETGRPRR<br>AAVSSFGISGTNAHVILEAPPEEPARDAAETRPEARSREAAEGRREAAGQGRTETGPGPSATPPEAPGRARPPVPWPLSGRDAGALR<br>DQIGRLRAHLDAAPADPEDVAHSLARRAVFRHRAVLLAAPQAPAGGSPRAVTGVARPGGTALLFSGQGSQRVGMGSELYETYPVFAE<br>SFDAVAEHTGLPLKDVVLGGTPDGLLDRTRYAQPALFAVEVSLFRLVRALGLDVRAVVGHSVGEIAAAHVAGVMSMADACRLVEARG<br>RLMDALPPGGAMVAVEVTEAEASAALAGLEDRVAVAAVNGPASTVLSGEEGAVLKLADAWRERGVRTHRLTVSHAFHSPLMEPMVDA<br>FREVVAGLDLHRPTLAGLPAEVVDPEYWVRHVRRPVRFADAVARAREAGAVRWLEVGPGGVLTALAQRIVPDTEEHVFAAALRTDRP<br>EPEALLVALSQVHVDGGTVDWSGLCAGGRLVDLPTYPFQRQHYWIEDQPLPPTAPRPGTAPSGTGTAAEGAAAAEVPLSERLARLTG<br>AERLAAVRELVLAEASETLGHTGTLITADRTRQELGFDSLTAIELRNRISRTLGVRLPPTLVFDHEDLGEIASFVDARLDDAATGRS<br>TGHGPLGEDGSGLLTELFREAAAAGRLDDAVTLTEAAARMRRTFTDAEDPAVRRTPVWFGRGPARPTVVCLPSFSAIAGVHVYARFA<br>DAFGDGWRVAALAHPGFVPGEPLPDSVDVLAELHARTVLDTVGADPFLLVGRSAGGWVAHEVAAVLERMGRAPDGVALLDTPARADD<br>PRGHAVMVGGMLERDSRLVTIDDYRLTAMGGYSRLFREWKPEPIAAATLLVHAATPYGADEARIASWDLPHQAVKVTGDHFTMLERH<br>SATTAEAVEQWSRSLLKLAAALEHHHHHHHH |

**Table S6.** Melting temperatures of the proteins composing the native and engineered VEMSs determined by thermal shift assay[6]

| Module(s) | PKS | Construct | $T_m$ in °C |
|---|---|---|---|
| M0-M1 | native VEMS | CS004.1 | 42.3±1.0 |
| M2 | native VEMS | CS005 | 40.0±0.0 |
| M0-M1 | DD-less VEMS | JH015 | 41.0±0.5 |
| M2 | DD-less VEMS | JH008 | 40.0±0.0 |
| M0-M1 | DEBS DD swap (α1-α3) | JH013 | 40.3±1.8 |
| M0-M1 | DEBS DD swap (α3) | JH014 | 41.5±0.0 |
| M2 | DEBS DD swap (α4) | JH005 | 40.3±0.3 |
| M0-M1 | SYNZIP-linked VEMS | CS002.2 | 42.3±0.3 |
| M2 | SYNZIP-linked VEMS | CS003 | 40.0±0.0 |
| M0-M1 | Spy-linked VEMS | JH017 | 40.3±0.3 |
| M2 | Spy-linked VEMS | JH010 | 40.3±0.3 |
| M0 | Split VEMS / Split SYNZIP-linked VEMS | LB047 | 44.3±0.3 |
| M1 | Split VEMS | CS008 | 39.3±0.3 |
| M1 | Split SYNZIP-linked VEMS | LB048 | 41.3±0.3 |

**Table S7.** Amino acid sequences used for the complex prediction of the native VemG:VemH docking interface (Figure 1) with ColabFold[1].

| Domain | Amino acid sequence |
|---|---|
| VemG $^{ACP}$DD | GIRAAEGTVFGALARLEASVDRDRPDAAARDRLVERLRALTARLTEGPAGGPSADDTSIGDRIGEATTVDQLFDLIDNELKGP |
| VemG $^{ACP}$DD | GIRAAEGTVFGALARLEASVDRDRPDAAARDRLVERLRALTARLTEGPAGGPSADDTSIGDRIGEATTVDQLFDLIDNELKGP |
| VemH $^{KS}$DD | MTGTEEKLVDYLKKVTAELQETRRQLRGALAASREP |
| VemH $^{KS}$DD | MTGTEEKLVDYLKKVTAELQETRRQLRGALAASREP |

**Table S8.** Amino acid sequences used for the complex prediction of the SYNZIP VemG-SZ3:SZ4-VemH docking interface (Figure S4) with ColabFold[1].

| Domain | Amino acid sequence |
|---|---|
| VemG ACP1-SZ3 | GPDRLRADLAPLTDDERERRLTDLVRGEIAAVAGFDGPHEIEAHRSMTDLGLDSVSAVDLSTRLGARTGLDLPASLAFDHPTST<br>AIARHLLAALGGSGGGSGNEVTTLENDAAFIENENAYLEKEIARLRKEKAALRNRLAHKK |
| VemG ACP1-SZ3 | GPDRLRADLAPLTDDERERRLTDLVRGEIAAVAGFDGPHEIEAHRSMTDLGLDSVSAVDLSTRLGARTGLDLPASLAFDHPTST<br>AIARHLLAALGGSGGGSGNEVTTLENDAAFIENENAYLEKEIARLRKEKAALRNRLAHKK |
| VemH SZ4-KS2 | MQKVAELKNRVAVKLNRNEQLKNKVEELKNRNAYLKNELATLENEVARLENDVAEGGSGGGSGKLEPIAIVGMACRYPGGVRTP<br>EALWRLVLDEQDAISGFPTNRGWDIDGIYHPDPDRPGTCYAREGGFLHDAALFDAEFFGVSPREAQAMDPQQRLLLETAWEAFE<br>RAGIDPTSLRGSDTGVFAGVVHHDYATARVPETLEPYLVTGLSGGVASGRIAYTFGFEGPAVTVDTACSSSLVALHQAAHALRS |

| Domain | Amino acid sequence |
| --- | --- |
| | GECELALAGGVTIMATPRAFLSFSRQRGLSPDGRCRAFGAGADGTGWAEGAGMLLVERLSDARRKGHPVLAVLRGSAVNQDGAS NGLSAPNGPSQQRVIRKALAHAGLAARDVDVVEGHGTGTKLGDPIEAQALLATYGQERDAGLPLHLGSMKSNVGHSQAAAGVGG VIKMIEAMRHGILPRTLHADEPTPHVDWSAGDIELLTRRRAWPETGRPRRAAVSSFGISGTNAHVILEAPP |
| VemH SZ4-KS2 | MQKVAELKNRVAVKLNRNEQLKNKVEELKNRNAYLKNELATLENEVARLENDVAEGGSGGGSGKLEPIAIVGMACRYPGGVRTP EALWRLVLDEQDAISGFPTNRGWDIDGIYHPDPDRPGTCYAREGGFLHDAALFDAEFFGVSPREAQAMDPQQRLLLETAWEAFE RAGIDPTSLRGSDTGVFAGVVHHDYATARVPETLEPYLVTGLSGGVASGRIAYTFGFEGPAVTVDTACSSSLVALHQAAHALRS GECELALAGGVTIMATPRAFLSFSRQRGLSPDGRCRAFGAGADGTGWAEGAGMLLVERLSDARRKGHPVLAVLRGSAVNQDGAS NGLSAPNGPSQQRVIRKALAHAGLAARDVDVVEGHGTGTKLGDPIEAQALLATYGQERDAGLPLHLGSMKSNVGHSQAAAGVGG VIKMIEAMRHGILPRTLHADEPTPHVDWSAGDIELLTRRRAWPETGRPRRAAVSSFGISGTNAHVILEAPP |

**Table S9.** Amino acid sequences used for the complex prediction of the DEBS2:DEBS3 docking interface (Figure S6) with ColabFold[1].

| Domain | Amino acid sequence |
| --- | --- |
| DEBS2 ACP4-$^{ACP}$DD | PNVVDRLAGRSESDQVAGLAELVRSHAAAVSGYGSADQLPERKAFKDLGFDSLAAVELRNRLGTATGVRLPSTLVFDHPTPLAV AEHLRDRLFAASPAVDIGDRLDELEKALEALSAEDGHDDVGQRLESLLRRWNSRRADAPSTSAISEDASDDELFSMLDQRFGGG EDL |
| DEBS2 $^{ACP}$DD | VDIGDRLDELEKALEALSAEDGHDDVGQRLESLLRRWNSRRADAPSTSAISEDASDDELFSMLDQRFGGGEDL |
| DEBS3 $^{KS}$DD-KS | MSGDNGMTEEKLRRYLKRTVTELDSVTARLREVEHRAGEPIAIVGMACRFPGDVDSPESFWEFVSGGGDAIAEAPADRGWEPDP DARLGGMLAAAGDFDAGFFGISPREALAMDPQQRIMLEISWEALERAGHDPVSLRGSATGVFTGVGTVDYGPRPDEAPDEVLGY VGTGTASSVASGRVAYCLGLEGPAMTVDTACSSGLTALHLAMESLRRDECGLALAGGVTVMSSPGAFTEFRSQGGLAADGRCKP FSKAADGFGLAEGAGVLVLQRLSAARREGRPVLAVLRGSAVNQDGASNGLTAPSGPAQQRVIRRALENAGVRAGDVDYVEAHGT GTRLGDPIEVHALLSTYGAERDPDDPLWIGSVKSNIGHTQAAAGVAGVMKAVLALRHGEMPRTLHFDEPSPQIEWDLGAVSVVS QARSWPAGERPRRAGVSSFGISGTNAHVIVEEAP |
| DEBS3 $^{KS}$DD-KS | MSGDNGMTEEKLRRYLKRTVTELDSVTARLREVEHRAGEPIAIVGMACRFPGDVDSPESFWEFVSGGGDAIAEAPADRGWEPDP DARLGGMLAAAGDFDAGFFGISPREALAMDPQQRIMLEISWEALERAGHDPVSLRGSATGVFTGVGTVDYGPRPDEAPDEVLGY VGTGTASSVASGRVAYCLGLEGPAMTVDTACSSGLTALHLAMESLRRDECGLALAGGVTVMSSPGAFTEFRSQGGLAADGRCKP FSKAADGFGLAEGAGVLVLQRLSAARREGRPVLAVLRGSAVNQDGASNGLTAPSGPAQQRVIRRALENAGVRAGDVDYVEAHGT GTRLGDPIEVHALLSTYGAERDPDDPLWIGSVKSNIGHTQAAAGVAGVMKAVLALRHGEMPRTLHFDEPSPQIEWDLGAVSVVS QARSWPAGERPRRAGVSSFGISGTNAHVIVEEAP |
